# Single-Cell and Spatial Methods for Multimodal Functional Glycan Profiling in Tissues

**DOI:** 10.64898/2026.02.23.707224

**Authors:** Ankit Basak, Carolina Ortiz-Cordero, Stephanie Pei Tung Yiu, Sidney Allison, Sarah Sweeting, Feng Shan, Yuzhou Evelyn Tong, Rajeev Chorghade, Alexxandra Sosa-Guir, Somkene D. Alakwe, Constantine Tzouanas, Bokai Zhu, Adele Gabba, Yao Yu Yeo, Wenrui Wui, Huaying Qiu, Yuzhou Chang, Cankun Wang, Eric J. Carpenter, Shreyas Gupta, Guilherme Meira Lima, Lindsay Parmelee, Philip Rock, Yi Zhang, Ratmir Derda, W Richard Burack, Qin Ma, Alex K. Shalek, Sizun Jiang, Laura Kiessling

## Abstract

Glycans regulate multiple physiological processes, including immune recognition and cancer progression. In disease, altered glycan landscapes are interpreted by human lectins. Functional glycan-lectin interactions are difficult to profile because glycans are not genome-encoded and their changes are poorly captured by existing multimodal methods. We present two platforms, single-cell outlining and transcriptome sequencing (scGOAT-seq) and GlycoScope, which use human lectins to enable functional glycan accessibility into single-cell and spatial multiomic measurements. ScGOAT-seq quantifies lectin-accessible glycan states with gene expression, while GlycoScope enables multiplexed in situ co-detection of glycans and proteins in tissues. Applying these approaches to immune cells, we identify stimulus-specific glycan remodeling and show that distinct Siglec-ligand-defined programs stratify immune activation states not captured by traditional methods; in follicular lymphoma, GlycoScope, resolves spatial glycan programs associated with malignant B cells and localized immune microenvironments. The presented methods provide a general framework for integrating functional glycan accessibility into single-cell and spatial multiomics.

## Introduction

Cell-surface glycans are essential regulators of cell communication and signaling, governing key biological processes such as immune recognition, pathogen entry, and cancer progression^1–6^. Their recognition by lectins, glycan-binding proteins, is central in coordinating both innate and adaptive immune responses. In cancer, glycosylation patterns are often profoundly altered^7^, producing tumor-associated glycan profiles that can drive immune evasion, metastasis, and resistance to therapy. As a result, deciphering how glycans are organized on cells in health and disease and how they are read by endogenous lectins is critical for understanding immune regulation and identifying biomarkers and therapeutic targets.

Recent single-cell multimodal profiling approaches – including scRNA-seq, CITE-seq, spatial proteomics, and spatial transcriptomics – have improved our ability to map cellular heterogeneity and spatial organization in tissues^8,9^. However, these approaches are not designed to capture relevant glycan–lectin interactions *in situ*, a critical axis of immune regulation. Glycan profiling remains challenging because glycan structures are not directly encoded in the genome, and mass spectrometry-based glycomics, while powerful, are not easily adaptable with simultaneous high-throughput single-cell or high-resolution spatial measurements. To address this gap, several groups have developed single-cell glycan profiling approaches using oligonucleotide-barcoded plant lectins or chemically modified antibodies. These methods have demonstrated that lectin-based barcodes can be read out together with RNA, and one of them, SUGAR-seq showed that specific glycan epitopes can be quantified at single-cell resolution^10,11^. These studies established feasibility but relied on non-human lectins or probed single glycan specificities per experiment; the underlying methods do not directly report how multiplexed human lectin receptors interpret glycan changes or extend into integrated spatial profiling of glycans and proteins.

Plant lectins are used widely as glycan sensors as they are invaluable tools for recognizing broad glycan classes, but human lectins are our endogenous glycan readers. Soluble human lectins operate in diverse settings, including in the blood and at mucosal surfaces. Membrane-bound lectins on immune cells survey their surroundings to regulate adhesion, signaling, and effector responses. Converting human lectins into molecular probes provides a direct, physiologically grounded way to assess cell-surface glycan states and how the immune system interprets them. For example, Siglecs, a family of sialic acid-binding immunoglobulin-type lectins, are expressed across various immune cell populations and mediate critical decisions in self- versus non-self-recognition, influencing inflammation and immune cell activation^12^. Calcium-dependent (C-type) lectins include members like DC-SIGN, a membrane-bound lectin expressed on macrophages and dendritic cells that binds high-mannose and Lewis-type antigens to mediate cellular adhesion, migration, signaling, and antigen presentation^13^; mannose-binding lectin (MBL), a soluble C-type lectin, recognizes glycans such as mannose and N-acetylglucosamine (GlcNAc), leading to the activation of complement pathways and modulation of inflammatory responses^14^; and, galectins, another class of soluble lectins, bind β-galactose-containing glycoconjugates and function in cell signaling, immune homeostasis, and tumor immunity^15^. Despite their biological importance, glycan–lectin interactions in human tissues remain poorly characterized, particularly at the single-cell and tissue resolutions.

To overcome these limitations, we sought to realize approaches that assess glycans using the receptors that detect them in human tissues. To this end, we developed two complementary platforms for profiling glycan heterogeneity in tissues that exploit the modular barcoding of human lectins. The first is single-cell glycan outlining and transcriptome sequencing (scGOAT-seq), which integrates human lectin-based glycan measurements with dissociated single-cell omic profiling, while the second is GlycoScope, which enables glycan-augmented spatial single-cell omics. By leveraging a customized panel of DNA-barcoded recombinant human lectins with immunological functions^15–17^, our methods overcome the limitations of conventional probes, enabling the detection of functional glycan recognition. Specifically, we define functional glycan states as lectin-accessible glycan features that reflect how endogenous receptors interpret the cell surface. Using scGOAT-seq across immune perturbations, we uncover stimulus-specific glycan remodeling programs that diverge between innate inflammatory and T cell receptor-driven activation; these include distinct Siglec-7-, Siglec-9-, and Siglec-15–defined glycan states that stratify immune phenotypes not resolved by transcriptomics alone. Functional perturbation by blocking endogenous human lectins further reveals direct modulation of immune activation and exhaustion-associated signaling programs, underscoring the critical roles of protein–glycan interactions in regulating immune cell states. Extending these measurements to tissues with GlycoScope, we identify spatially organized glycan programs in follicular lymphoma that map to malignant B cell populations and structured immune microenvironments. In summary, our work provides a robust framework for integrating functional glycan profiling with existing single-cell and spatial methods to enhance cell-state stratification in tissues towards the discovery of multimodal biomarkers and therapeutic targets in cancer and beyond.

## Results

### Design and validation of a recombinant lectin panel for single-cell glycan profiling

To generate a panel of recombinant lectins compatible with multiplex single-cell glycan profiling, we curated a panel of 7 human recombinant lectins engineered to bear a biotin label (Fig. S1A). Our panel includes inhibitory lectins (e.g., Siglec-7, Siglec-9), others that deliver context-dependent signals (Siglec-15, Galectin-8, Galectin-9 and DC-SIGN), and the complement-recruiting MBL. This set recognizes major mammalian glycan classes, including sialylated glycans, N-acetyllactosamine (LacNAc)/poly-LacNAc, mannose-rich structures, and fucosylated motifs, providing a functionally informative panel for cellular glycan profiling.

To ensure the binding preferences of our tagged lectins were unchanged, we profiled their specificity using a liquid glycan array (LiGA; Fig. 1A)18. Concanavalin A (ConA) and UEA-I were included as assay controls to benchmark high-mannose and α-fucose recognition, respectively. Hierarchical clustering revealed three coherent specificity modules aligned with known glycan–lectin biology12–15. Siglec-7, Siglec-9, and Siglec-15 bound sialic acid-containing glycans. Moreover, Siglec-7 and Siglec-9 showed the strongest signal for α(2,3)- and α(2,6)-sialylated LacNAc and glycolipid-like structures. Both also showed moderate binding to a subset of neutral LacNAc and extended poly-LacNAc glycans, a pattern consistent with multivalent display. Siglec-15 displayed a more restricted footprint with a bias toward α(2,6)-linked sialic acid. Galectin-8 and Galectin-9 clustered together and showed maximal binding to LacNAc and extended poly-LacNAc structures, with markedly reduced recognition of their sialylated counterparts. A third module captured pathogen-associated glycan recognition. MBL bound strongest to oligomannose structures but also showed moderate engagement of GlcNAc-rich glycans. DC-SIGN overlapped with this mannose-binding pattern and additionally recognized fucosylated motifs.

**Fig 1.**
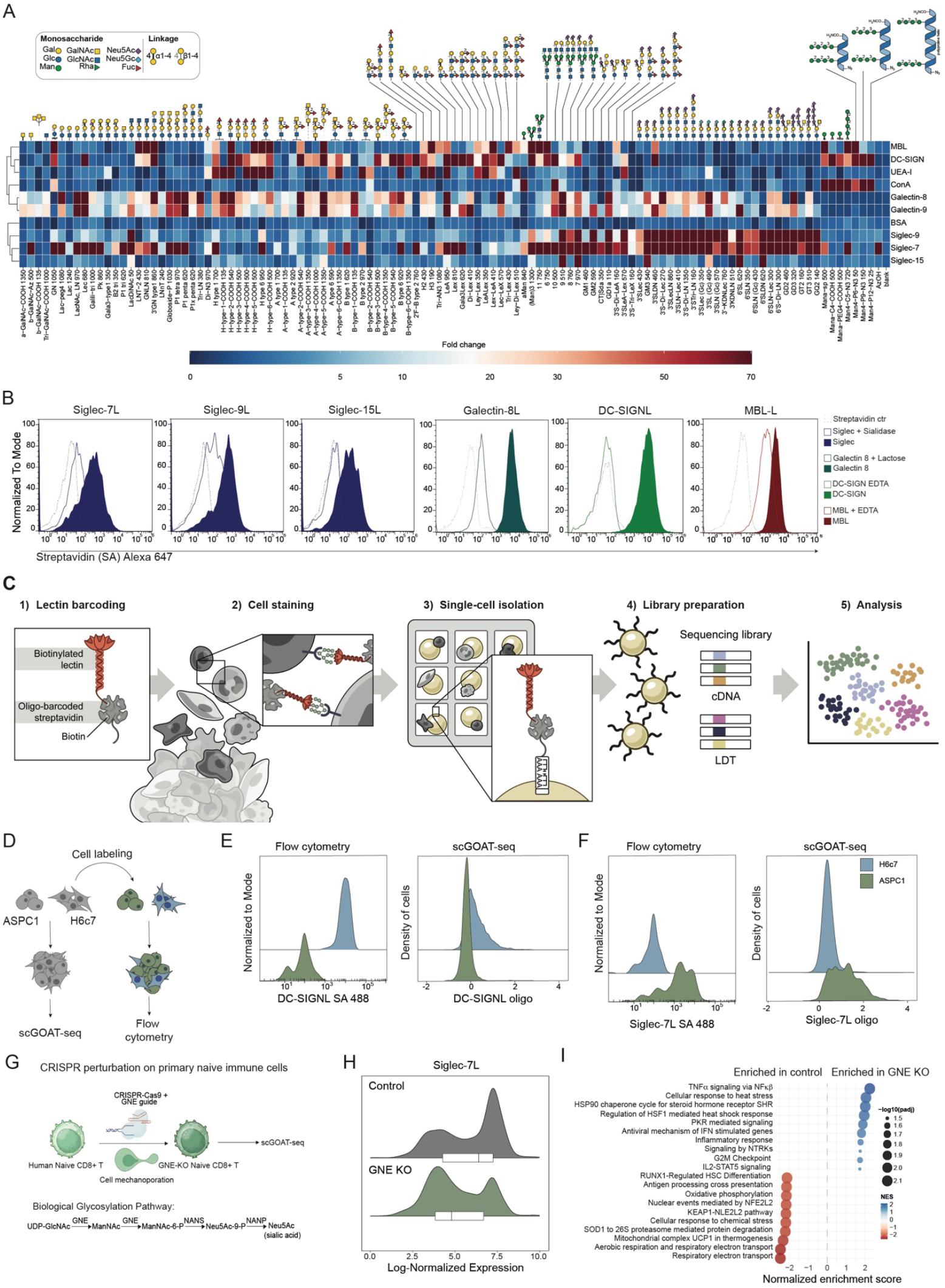
scGOAT-seq enables multimodal single-cell detection of glycan abundance using human recombinant lectins alongside gene expression. **(A)** Hierarchically clustered heatmap of lectin binding signals to LiGA. Columns correspond to glycans labeled with the density that maximized discriminatory behavior across lectins. Binding intensity is shown as fold change where red indicates high enrichment and blue indicates low or absent binding. Scale bar is quadratic with fold changes 0–70. **(B)** Representative histograms of flow cytometry analysis confirming lectin binding, with respective negative controls: sialidase for Siglec-7, -9, and –15 in ASPC1 cells, lactose for Galectin-8 in Toledo cells, EDTA for DC-SIGN in H6c7 cells, and MBL in A549. **(C)** Schematic of single-cell glycan outlining and transcriptome sequencing (scGOAT-seq) assay using barcoded recombinant human lectins. **(D)** Overview of the benchmarking experiment with an artificial pancreatic tumor dissociation model, consisting of cancer (ASPC1) and non-cancer (H6c7) pancreatic cell lines, deconvoluted using flow cytometry (dye-based) and scGOAT-seq (transcriptomic reference map-based), followed by glycan profiling comparison. **(E)** Representative histograms showing DC-SIGNL and **(F)** Siglec-7L abundance per cell line in the artificial pancreatic tumor dissociation model analyzed by both flow cytometry and scGOAT-seq. **(G)** Schematic representation of GNE gene CRISPR KO experimental setup on human naive CD8+ T cells and the role of GNE in sialic acid biosynthesis pathway. **(H)** Representative scGOAT-seq histograms showing Siglec-7L in GNE-KO versus control CD8+ T cells. **(I)** Gene set enrichment analysis comparing naïve versus GNE-KO CD8+ T cells.

To confirm binding specificity in a cellular context, lectins were assayed by flow cytometry following cell-surface staining with biotinylated lectins and detection using fluorescently conjugated streptavidin. Binding specificity was evaluated using orthogonal perturbations, including enzymatic removal of target glycans (sialidase treatment), chelation of calcium for C-type lectins (EDTA), or competition with soluble glycans (lactose). Each perturbation resulted in a consistent reduction in lectin binding compared to untreated controls (Fig. 1B). Collectively, these results demonstrate that each lectin retains its expected biochemical behavior and provides a validated framework for their downstream use as glycan-recognition probes.

Finally, we tested whether multiplexed staining caused cross-interference. Each lectin was profiled alone and in pairwise combinations using peripheral blood mononuclear cells (PBMCs). Similarity between single-lectin and paired-lectin fluorescence distributions was quantified using the Bhattacharyya overlap coefficient. Across the human lectin panel, overlap values remained high, with little evidence of competitive binding or altered specificity. In contrast, several plant lectins (ConA, PHA-L) showed reduced overlap, likely reflecting lectin–lectin interactions or steric masking. These data indicate that co-incubation preserves the fidelity of human lectin binding and supports the use of the panel for multiplexed glycan detection (Fig. S1B). Together, these analyses demonstrate that human lectins maintain independent, non-interfering binding behavior under multiplex conditions.

### scGOAT-seq enables multimodal detection of functional glycan patterns and gene expression programs

To simultaneously measure single-cell glycan abundance and gene expression, we equipped recombinant human lectins with poly(A)-tailed DNA barcodes containing PCR handles, cell barcodes and unique molecular identifier sequences (UMIs)^19^, enabling their capture alongside mRNA in standard poly(A)-based single-cell transcriptomic workflows (**Fig. 1C**). Direct covalent coupling of DNA barcodes impaired binding for several lectins (**Fig. S1C**). We therefore adopted a biotin–streptavidin strategy ^20–22^, generating lectin–barcode conjugates in ∼30 min and enabling flexible panel design (**Fig. S1D**). We evaluated our conjugation strategy, using flow cytometry to compare the binding performance of streptavidin conjugated to either a DNA oligonucleotide or a fluorescent dye. As a representative example, we assessed DC-SIGN, a calcium-dependent (C-type) lectin, and found that the oligonucleotide-conjugated streptavidin exhibited binding comparable to its fluorophore labeled counterpart. This interaction was calcium-dependent, confirming that oligo conjugation does not impact lectin binding (**Fig. S1E**).

Sialylated glycans in pancreatic adenocarcinoma (PDAC) cells have been reported to drive immunosuppression through binding to Siglec-7 and -9 on immune cells ^23^. To test the robustness of scGOAT-seq in an *in vitro* cancer model, we mixed a human pancreatic cancer cell line, ASPC1, known to have distinct glycan profile^24^, with a non-malignant pancreatic epithelial cell line (H6c7; **Fig. 1D**). To evaluate the multiplexing capability of scGOAT-seq, we used four lectins (Siglec-7, Siglec-9, Siglec-15 and DC-SIGN) for profiling (**Fig. 1B**).

We quantified lectin binding by flow cytometry using cell-tracer dyes to distinguish cell lines and performed scGOAT-seq on an unstained mixed-cell population using Seq-Well S^3^ platform (**Fig. S1F**). To assign cell identities in the scGOAT-seq dataset, we mapped cells to a curated reference transcriptomic atlas for the PDAC model. We observed high correlation in gene expression signals between the atlas and cells treated with lectin (**Fig. S1G-H**), indicating that lectin staining did not substantially perturb transcriptional state. ScGOAT-seq reliably detected Siglec-accesible ligands (L) and DC-SIGNL in each cell type in close agreement with flow cytometry measurements. Both methods identified the highest binding to Siglec-L in ASPC1 cells and the lowest in H6C7, while DC-SIGNL binding was higher in H6C7 compared to ASPC1 (**Fig. 1E-F and Fig. S1I**). To further validate the specificity of scGOAT-seq, we treated our multicellular model with sialidase to enzymatically remove sialic acid epitopes. As expected, the level of different Siglec binding glycans in ASPC1 cells was reduced (**Fig. S1J**). This consistency underscores the reliability of scGOAT-seq for glycan profiling.

To examine how glycosylation shapes immune cell behavior, we downregulated sialic acid biosynthesis in primary human naïve CD8+ T cells, which display high levels of cell-surface sialic acids^25^. Using a mecha-noporation-based Cas9/sgRNA delivery system that minimizes activation of resting lymphocytes, we knocked out *GNE*, which encodes the rate-limiting enzyme for Neu5Ac biosynthesis from UDP-GlcNAc. GNE has been implicated as a determinant of Siglec-7 ligand display in genome-wide CRISPRi screens^26^ (**Fig. 1G**). We profiled *GNE*-KO and scrambled-control CD8+ T cells with scGOAT-seq (**Fig. 1H**). Loss of *GNE* resulted in a marked reduction in Siglec-7L binding relative to controls. Gene-set enrichment analysis (GSEA) showed that *GNE*-KO cells upregulated TNFα signaling, interferon responses, inflammatory pathways, and IL2-STAT5-induced signaling (**Fig. 1I**). These results support the utility of scGOAT-seq as a platform for linking perturbations of glycan biosynthesis to lectin-accessible glycan states and associated transcriptional programs at single-cell resolution.

Collectively, our benchmarking results demonstrate that scGOAT-seq yields glycan-binding signals closely aligned with flow cytometry and captures glycan changes induced by enzymatic and genetic perturbations in both cell lines and primary immune cells. Thus, this method enables high-resolution glycan profiling alongside single-cell gene expression, with integrability across a wide range of single-cell RNA-seq platforms.

### scGOAT-seq identifies novel glycan-defined cell states associated with immune activation in peripheral blood

The sialic acid-Siglec axis is an emerging glyco-immune pathway targeted by therapeutic agents that disrupt Siglec interactions, particularly in the context of cancer therapy^27,28^. These approaches are designed to block engagement between Siglec receptors on immune cells and sialylated ligands on tumor cells. However, how distinct Siglecs, and other lectin-mediated recognition pathways, respond to different immuno-modulatory cues remains incompletely defined. In addition, the transcriptional programs associated with glycan remodeling are poorly characterized in multi-cellular immune settings, in which different cell types express distinct lectin repertoires and can therefore interpret shared glycan landscapes in context-dependent ways.

To investigate these effects, we performed scGOAT-seq on PBMCs with either (i) bacterial lipopolysaccharide (LPS; 4 h) to stimulate innate immune responses^29^ or (ii) anti-CD3/CD28 (24 h) to induce T cell receptor (TCR) activation^30^ (**Fig. 2A**). Cells were then profiled by scGOAT-seq on the 10X Genomics 5’ platform using our curated human lectin panel. Across donors and conditions, we identified 8 different broad immune subtypes across 7,247 cells, including CD4+ and CD8+ T cells, NK cells, monocytes and B cells, using Azimuth-based reference mapping^31^ and marker genes (**Fig. 2B-C and Fig. S2A**).

**Fig 2.**
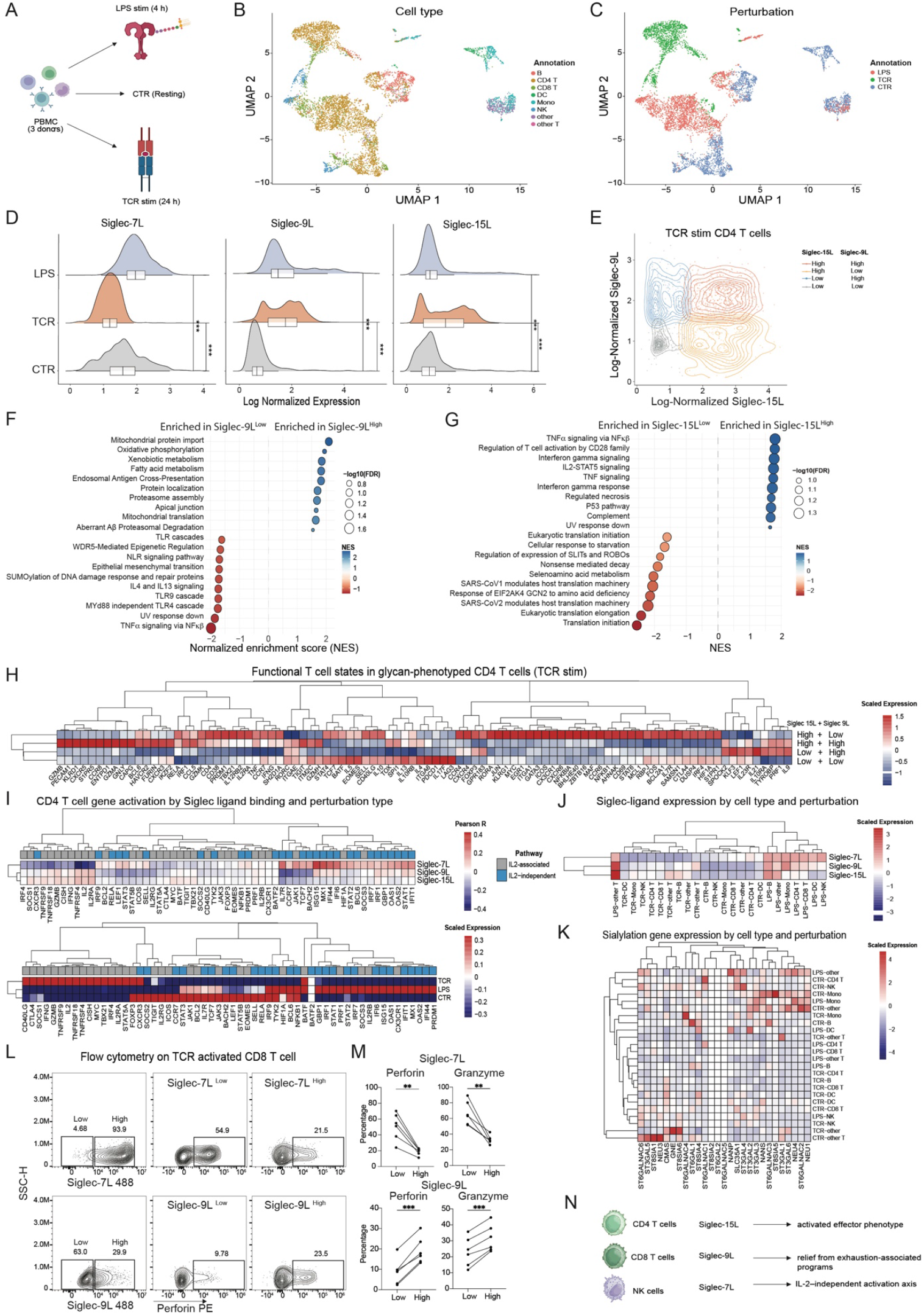
scGOAT-seq identifies novel glycan-defined cell states associated with immune perturbations. **(A)** Schematic figure showing perturbations of PBMCs with LPS stimulation and anti-CD3/anti-CD28 antibodies for TCR stimulation. **(B, C)** UMAP projections of different PBMC cells by cell type and perturbation. **(D)** Ridge plots depicting log-normalized expression of lectin-binding ligands for Siglec-7L, Siglec-9L and Siglec-15L in CD4+ T cells across perturbations. **(E)** Scatter density plot showing different populations of TCR stimulated CD4+ T cells binned by expression levels of Siglec-9L and Siglec-15L. **(F-G)** Gene set enrichment analysis of TCR stimulated CD4+ T cells by expression levels of Siglec-9L and Siglec-15L respectively. **(H)** Heatmap of key functional CD4+ T cell gene expression across TCR stimulated CD4+ T cells grouped by expression levels of Siglec-9L and Siglec-15L. **(I)** Heatmap showing Pearson correlation of functional CD4+ T cell activation genes with Siglec-7L, Siglec-9L and Siglec-15L (top) and their expression across CD4+ T cells in different perturbations (bottom). **(J)** Expression and clustering of Siglec-7L, Siglec-9L and Siglec-15L across different cell type and conditions. **(K)** Expression and clustering of all sialylation-related genes across different cell types and conditions. **(L)** Spectral flow cytometry gating of CD8+ T cells after TCR activation, with independent stratification by Siglec-7 or Siglec-9 ligand binding and assessment of perforin expression within each Siglec-defined population. **(M)** Frequencies of perforin- and granzyme-expressing CD8+ T cells following TCR activation, stratified by Siglec-7 or Siglec-9 ligand binding. **(N)** Schematic representation of glycan states associated with Siglec-7L, Siglec-9L and Siglec-15L across T and NK cells. NES: Normalized Enrichment Score. **P<0.01, ***P<0.001.

Immune stimulation induced distinct, perturbation- and cell-type–dependent changes in lectin binding profiles. For example, Siglec-7L binding was increased upon LPS stimulation but decreased after TCR stimulation. By contrast, Siglec-9L increased under both perturbations, whereas Siglec-15L was selectively upregulated following TCR stimulation and showed minimal change after LPS (**Fig. 2D and Fig. S2B-C**). Other lectin-accessible features, including DC-SIGNL and Galectin-8L binding increased across both activation conditions (**Fig. S2C**). Based on these divergent binding patterns within the same Siglec family, we hypothesized that binding of different Siglecs may be associated with different cellular functional transcriptomic states.

To investigate the association of different Siglecs with cellular states, we focused on CD4+ T cells, which exhibited bimodal populations identified by Siglec-9L and Siglec-15L after TCR stimulation. Cells were stratified into Siglec-ligand high and low populations based on the geometric mean of their expression level, and we performed GSEA to identify functional pathways associated with each glycan-defined subset (**Fig. 2E**). Siglec-9L^High^ CD4+ T cells were enriched in metabolic and respiratory pathways like xenobiotic metabolism, fatty acid metabolism, mitochondrial protein translation and import, whereas Siglec-9L^Low^ CD4+ T cells were enriched in activation-associated pathways like TNFα signaling, TLR9 downstream cascades and IL4/IL13 signaling pathways (**Fig. 2F**). Interestingly, Siglec-15L^High^ CD4+ T cells were enriched in activation pathways like TNFα signaling, CD28 co-stimulation pathway, IFN-γ pathway and IL2-STAT5 signaling (**Fig. 2G**). These results suggested that Siglec-9L^High^ CD4+ T cells were associated with an immuno-metabolic state whereas Siglec-15L^High^ CD4+ T cells were associated with an immune-activated state.

To further investigate how Siglec-9L/Siglec-15L relate to CD4+ T cell states, we compared gene expression levels across CD4+ T cell subsets stratified by their Siglec-ligand levels (**Fig. 2H**). Siglec-15L^High^/Siglec-9L^High^ cells exhibited a gene expression profile associated with cytotoxicity and inflammation (*GZMB, GNLY, IFNG, TNF*) and upregulated inhibitory receptors (*HAVCR2, TIGIT, TOX*), consistent with a highly activated effector phenotype. In contrast, Siglec-15L^High^/Siglec-9L^Low^ cells expressed higher levels of TCR activation genes (*JUN, FOS, EGR1, NFKBIA, NFKB1*), migratory markers (*CXCR4, CX3CR1,CXCR6*), co-stimulatory receptors (*ICOS, CD69, CTLA4*), and memory-associated genes (*S1PR1, IL7R*), indicating a transitional early-activation state. Siglec-15L^Low^/Siglec-9L^High^ cells exhibited minimal cytotoxic genes and expressed memory/stem-like genes (*LEF1, SPOCK2, KLF3*), while Siglec-15L^Low^/Siglec-9L^Low^ CD4+ T cells exhibited central memory features (*CCR7, SELL, TCG7, BATF*) and immune checkpoint expression (*PDCD1, LAG3, TIGIT, TOX*). Overall, these patterns show that Siglec-15L and Siglec-9L levels resolve distinct CD4+ T cell phenotypes across an activation continuum. Similarly, CD8+ T cells and NK cells showed distinct cell states associated with Siglec-9L/Siglec-15L phenotypes, with Siglec-15L^High^ states associated with more activated phenotypes (**Fig. S2C-D**).

Pathway level analysis further highlighted the difference in the Siglec axes. The prescence of Siglec-7L was negatively associated with the expression of IL-2-dependent activation genes, including *IL2, IL2RA*, and *IFN*`γ`. In contrast, these genes were positively correlated with detection of Siglec-9 and Siglec-15 binding. Conversely, Siglec-7 interactions were positively associated with IL-2-independent activation and memory-associated pathways, including *IL7R, CCR7, TCF7*, and *BACH2* – genes which were negatively correlated with the binding of Siglec-9 and Siglec-15 (**Fig. 2I**). The transcriptional patterns mirrored the perturbation context: IL-2-associated programs were strongly induced by TCR stimulation, whereas IL-2-independent pathways were preferentially upregulated following LPS treatment. Similar correlations between Siglec-7L and IL-2-associated activation pathways were observed in CD8+ T cells (**Fig. S2E**). These results highlight that Siglec ligands report on discrete activation states rather than a single inhibitory program, suggesting that Siglec-targeted therapies may differentially affect distinct arms of T cell activation depending on which Siglec axis is engaged.

Next, we examined whether Siglec-ligand profiles could classify perturbation states. Joint clustering of populations with enhanced binding to Siglec-7, Siglec-9, and Siglec-15 grouped all cells by activation condition rather than lineage, indicating that these three ligands are strong predictors of PBMC activation state (**Fig. 2J**). Broader lectin panels, including galectins and C-type lectins, did not improve clustering (**Fig. S2F**). In contrast, gene expression of all 27 genes associated with sialylation pathways (sialic acid biosynthesis, sialyl transferases and neuraminidases) showed distinct patterns across cell types but did not cluster the perturbations (**Fig. 2K**) with similar outcomes from analyzing O-glycosylation associated genes (**Fig. S2G)**.

To validate the association between Siglec-accessible ligand states and cytotoxic programs, we performed spectral flow cytometry on TCR-stimulated T cells and focused on CD8+ T cells. Cells were stained with Siglec probes alongside surface markers and intracellular cytotoxic effectors, including perforin (*PRF1*) and granzyme B (*GZMB*). For each Siglec, CD8+ T cells were stratified into Siglec-ligand–positive (Siglec-L^+^) and Siglec-ligand–negative (Siglec-L^−^) populations based on probe binding. Siglec-7L^−^ CD8+ T cells exhibited more than a two-fold higher frequency of perforin-positive cells compared to Siglec-7L^+^ cells. In contrast, Siglec-9L^+^ CD8+ T cells showed a higher proportion of perforin-positive cells than their Siglec-9L^−^ counterparts, consistent with transcriptomic trends observed by scGOAT-seq (**Fig. 2L and S3A-B**). These patterns were reproducible across six independent donors and were observed for both perforin and granzyme B, with significant differences in the fraction of effector-positive cells between Siglec-L^+^ and Siglec-L^−^ populations and opposing trends across distinct Siglecs (**Fig. 2M**). This suggests that binding of different Siglecs may be associated with distinct CD8+ T cell activation states both at transcriptomic and proteomic levels.

Together, these data indicate that Siglec-ligands are predictive of immune perturbation states of PBMCs and that different Siglec-ligand profiles are associated with distinct immune activation states (**Fig. 2N**).

### scGOAT-seq identifies immune activation changes associated with blocking lectin-mediated glycan regulation in peripheral blood

The stimulation-dependent shifts in Siglec-ligand accessibility observed under LPS and TCR activation suggest that changes in lectin binding reflect functional immune phenotypes mediated by Siglec–sialic acid recognition. Because monocytes and NK cells express the Siglec-7 and Siglec-9 receptors^32^, we reasoned that blocking these interactions would modulate immune activation trajectories upon stimulation. To test this, we repeated the LPS stimulation and scGOAT-seq workflow and included blocking antibodies against Siglec-7 and Siglec-9^33^ (**Fig. 3A**). Across donors and conditions, we again annotated 8 broad immune populations among 7,286 cells (**Fig. 3B-C, and Fig. S4A**).

**Fig 3.**
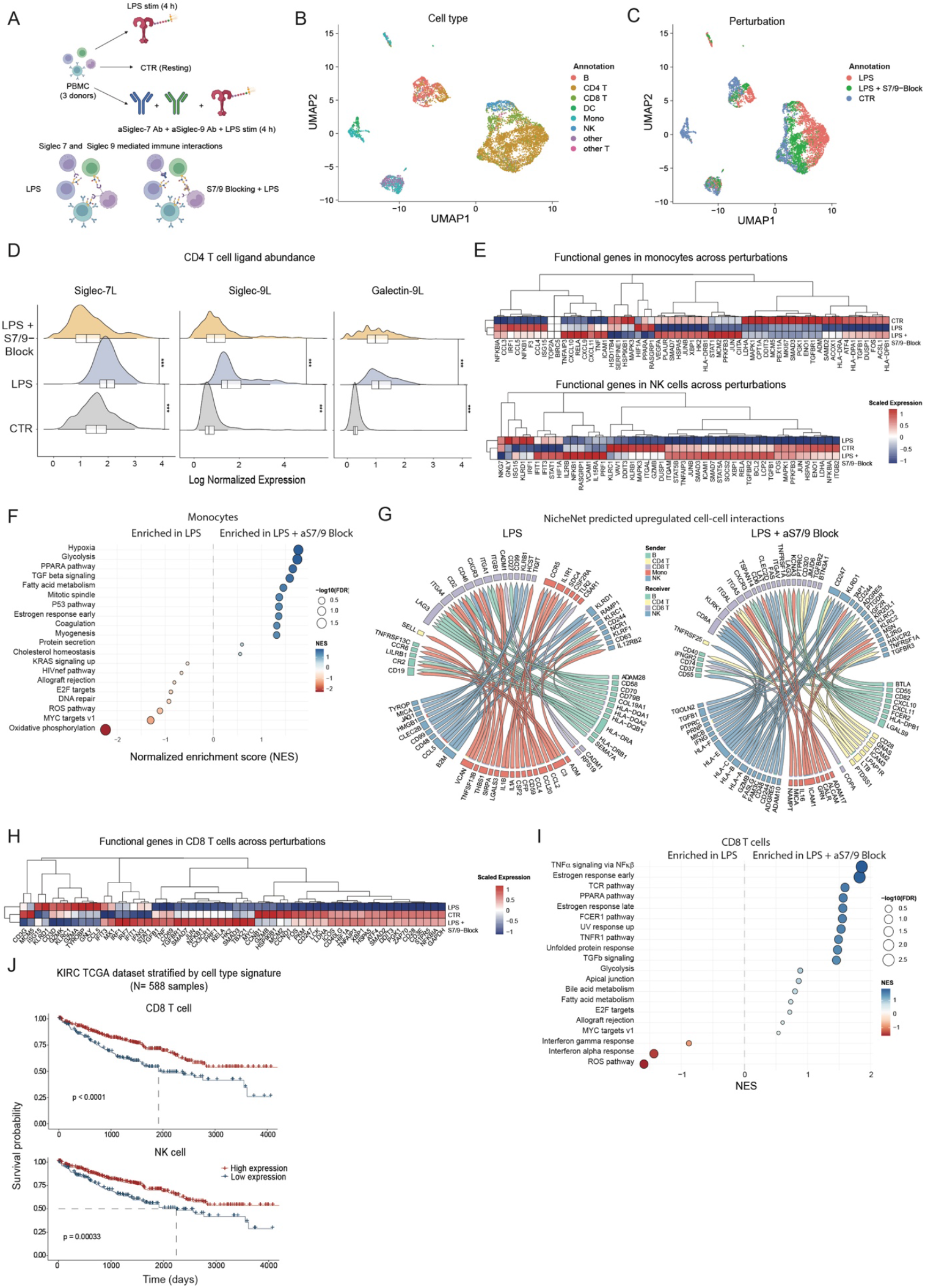
scGOAT-seq identifies immune activation changes associated with blocking Siglec-mediated glycan recognition. **(A)** Schematic figure showing perturbations of PBMCs with LPS stimulation and LPS stimulation with anti-Siglec7 and anti-Siglec9 blocking antibodies. **(B-C)** UMAP projections of different PBMC cells by cell type and perturbation. **(D)** Ridge plots depicting log-normalized expression of lectin-binding ligands for Siglec-7L, Siglec-9L and Galectin-9L across cell types and perturbations. **(E)** Heatmap of key functional monocyte (top) and NK cells (bottom) gene expression across perturbations. **(F)** GSEA analyses across monocytes between LPS stimulation with anti-Siglec7 and anti-Siglec9 blocking antibodies compared to LPS treatment alone. **(G)** NicheNet circus plots depicting top 100 prioritized potential cell-cell interactions in LPS treatment alone and LPS stimulation with anti-Siglec7 and anti-Siglec9 blocking antibodies. **(H)** Heatmap of key functional CD8+ T gene expression across perturbations. **(I)** GSEA analyses across CD8+ T cells between LPS stimulation with anti-Siglec7 and anti-Siglec9 blocking antibodies compared to LPS treatment alone. **(J)** Kaplan-Meier curves showing stratification of KIRC TCGA patient data by CD8+ T cell signature and NK cell signature induced by Siglec-blocking compared to LPS alone and stratified by top and bottom 30^th^ quantile of patient survival data. NES: Normalized Enrichment Score.

As expected, Siglec-7L binding induced by LPS was abolished in the presence of Siglec-7/9 blockade. Siglec-9L binding was reduced relative to LPS alone, although it remained higher than in resting PBMCs (**Fig. 3D**). Blocking Siglec receptors during LPS stimulation also broadly altered the binding of other lectins, including reduced binding of DC-SIGN, Galectin-8, MBL, and Siglec-15 (**Fig. S4B**). In contrast, Galectin-9 showed a distinct, cell type-dependent interaction pattern with binding decreases in T cells but increases in monocytes, DCs and NK cells (**Fig. 3D and Fig. S4B**). Although perturbations induced modest changes in the expression of O-glycosylation–related genes, these transcriptional effects did not recapitulate the observed lectin-binding patterns (**Fig. S4C**). Together, these results support a model in which LPS-driven glycan remodeling engages endogenous Siglec-7 and Siglec-9, and perturbation of these pathways propagates cell-type–specific rewiring of lectin–glycan interactions across the immune compartment.

We next examined transcriptional changes in monocytes and NK cells, two populations expected to respond directly to the Siglec blockade because they express Siglec-7 and Siglec-9^32^. In monocytes, Siglec blocking rewired inflammatory and metabolic programs, inducing cytokines (*CXCL9, CXCL10, TNF*), NFKB pathway genes (*RELA, JUN, JUNB, STAT1*), adhesion genes (*ICAM1, PLAUR*), metabolic genes (*VEG-FAM PFKFB3, HK2*), unfolded protein response genes (*XBP1, HSPA5*) and class II MHC genes (*HLA-DRB1, CIITA*). NK cells showed a parallel shift, with increased NFKB genes (*RELA, JUN, NFKBIA, TNFAIP3*), cytotoxic genes (*GZMB, PRF1*), interferon genes (*IFIT1, IFIT3, STAT1, IRF1, STAT5A*) and genes associated with heightened cytokine sensitivity (*IL15RA,IL2RB*) (**Fig. 3E**). GSEA analysis on monocytes revealed enrichment in hypoxia, glycolysis, PPARA pathway, TGFβ signaling and fatty acid metabolism, alongside reduced oxidative phosphorylation and ROS pathways (**Fig. 3F**). Overall, these findings suggest disrupting Siglec-7/9 engagement amplifies immune activation and metabolism pathways in monocyte and NK cell compartments under LPS stimulation.

To determine whether disrupting Siglec–sialic acid recognition alters intercellular immune communication, we used NicheNet^34^ to infer ligand–receptor interactions linked to downstream transcriptional responses. Analysis of the top predicted interactions revealed reduced monocyte-centered signaling, with monocytes showing decreased activity both as signal senders and receivers, alongside increased interaction activity involving CD4+ T cells as signal senders and NK cells as both senders and receivers. Under LPS stimulation alone, the interaction landscape was dominated by monocyte-derived inflammatory signals, including *CCL2, CCL3*, and *IL1B*. In contrast, Siglec-7/9 blockade shifted the network toward enhanced IFNγ- and cytotoxic-associated signaling (*IFNG, GZMB*), as well as increased co-stimulatory pathways involving *CD28, FAS*, and *HAVCR2* (**Fig. 3G**). Together, these changes indicate that disrupting Siglec-7/9 engagement reprograms immune communication networks toward amplified inflammatory signaling and NK cell– driven activation.

Gene expression changes were also examined in CD8+ T cells. While not expected to interact directly with anti-Siglec-7/9 antibodies, these cells can give rise to secondary responses through altered cell–cell signaling^32^. CD8+ T cells had heightened expression of inter-feron-induced genes (*IFIT3, IFIT1, MX1, STAT*1) and *TNF*, consistent with indirect activation driven by interferon-mediated communication from cell-cell interaction changes (**Fig. 3H**). GSEA analysis of CD8+ T cells from PBMCs treated with LPS with Siglec blocking showed enrichment of TNFα signaling, and the TCR and TNFR1 pathways, along with reduced IFN-α responses (**Fig. 3I**). Similar pathway activation, including PI3K-AKT, IL2-STAT5, hypoxia and PYK2 signaling, occurred in CD4+ T cells upon Siglec blocking (**Fig. S4 D-E**). Overall, these results suggest blocking Siglec-sialic acid interactions in PBMCs can drive both direct immune activation and secondary immune activation through altered cell-cell communication and signaling networks.

To probe the potential clinical relevance of the Siglec-blockade–associated immune programs, we analyzed bulk RNA-seq data from The Cancer Genome Atlas (TCGA) with matched survival information. We focused on kidney renal clear cell carcinoma (KIRC), a tumor type in which sialylation-associated signatures can stratify patient outcomes^28^, and restricted the analysis to patients without prior neoadjuvant therapy. We performed module-based gene scoring with the CD8+ T and NK cell signatures derived from genes upregulated and downregulated by LPS + Siglec blocking relative to LPS alone. Both signatures significantly stratified patient survival, with higher expression associated with better survival outcomes (**Fig. 3J**). Together, these results suggest that the immune activation programs revealed by Siglec-7/9 blockade reflect biologically meaningful anti-tumor immune states that are associated with favorable prognosis in kidney cancer.

### GlycoScope enables robust single-cell multiplex spatial co-detection of glycans and proteins

ScGOAT-seq provides a powerful framework to quantify glycan binding and associated transcriptional programs at single-cell resolution, offering mechanistic insights that cannot be obtained from tissue sections alone. However, many glycan-mediated processes are fundamentally shaped by their spatial organization, in which glycans, lectin receptors, and neighboring cells are positioned within native microanatomical niches. Current spatial multiomic platforms can resolve proteins and cell states *in situ*, providing critical insight into cellular interactions and tissue architecture that underlie spatially regulated biological processes^35^. However, they do not provide functional glycan-binding readouts, leaving a critical gap in the field. To address this unmet need and to complement the molecular insights gained from scGOAT-seq, we developed GlycoScope, a complementary multimodal spatial technology that leverages our barcoding strategy to equip recombinant human lectins with oligonucleotide tags compatible with CODEX-based proteomic imaging. This configuration enables single-cell, *in situ* detection of both glycans and proteins within intact tissue architecture (**Fig. 4A**), extending lectin binding into the spatial domain and providing an integrated view of glycan biology in its native context.

**Fig 4.**
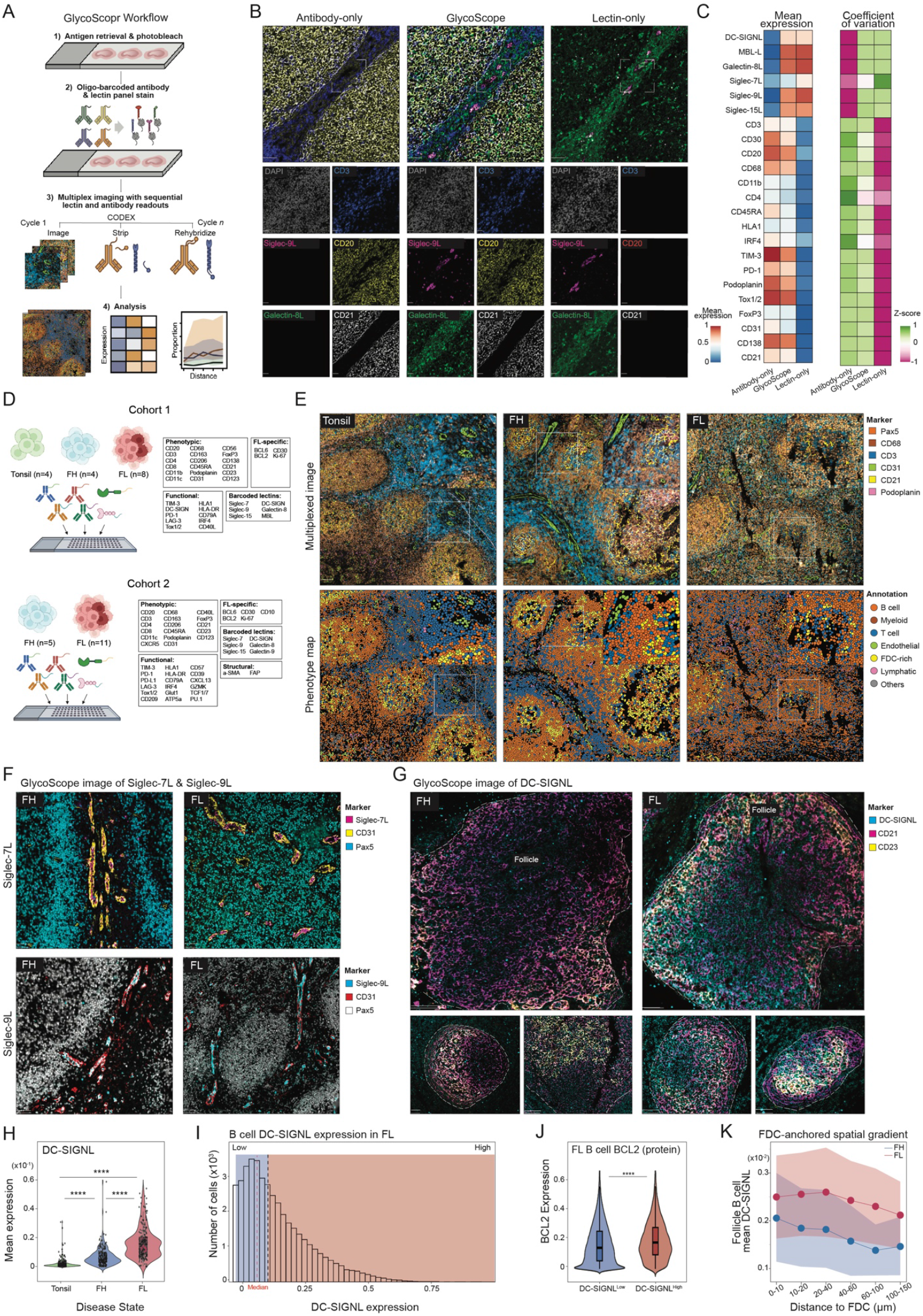
GlycoScope enables multiplex spatial glycan-protein detection and, together with scGOAT-seq, recapitulates mannosylated-BCR biology in follicular lymphoma. **(A)** Schematic workflow of GlycoScope, illustrating the sequential staining of antibody and recombinant lectin panels. **(B)** Representative multiplexed images showing nuclei (DAPI), Siglec-9L, Siglec-15L, Galectin-8L, T cells (CD3), B cells (CD20), and follicular dendritic cells (CD21) across GlycoScope, Antibody-only, and Lectin-only conditions of tonsil tissues (scale bar: 50µm, 10µm (inset)). All individual images across these three conditions are in **Fig. S4B. (C)** Heatmap showing the 0,1 normalization of mean expression (left) and z-score of the coefficient of variation (right) of all glycan and protein markers obtained from GlycoScope, Antibody-only, and Lectin-only conditions. **(D)** Schematic overview of the GlycoScope study cohorts. Cohort 1 consists of an FFPE tissue microarray (TMA) containing four tonsil, four follicular hyperplasia (FH), and eight follicular lymphoma (FL) cores, profiled using a 32-plex protein panel and six oligo-conjugated lectins (Siglec-7, Siglec-9, Siglec-15, MBL, DC-SIGN, and Galectin-8). Cohort 2 consists of an FFPE TMA containing 11 FL and 5 FH samples, profiled using a 42-plex protein panel and six oligo-conjugated lectins (Siglec-7, Siglec-9, Siglec-15, DC-SIGN, Galectin-8, and Galectin-9). **(E)** Representative GlycoScope multiplexed images (top) showing nuclei (DAPI), B/tumor cells (Pax5), macro-phages (CD68), T cells (CD3), endothelial cells (CD31), FDC-enriched region (CD21) and lymphatic (Podoplanin), as well as the corresponding phenotype maps (bottom) of FL, FH and tonsil tissues (scale bar: 100µm, 20µm (inset)). All individual phenotype maps for each tissue sample core are in **Fig. S5A. (F)** Top: Representative GlycoScope multiplexed images of FL and FH showing Siglec-7L (magenta), CD31 (yellow), and PAX5 (cyan). Bottom: Representative GlycoScope multiplexed images of FL and FH showing Siglec-9L (cyan), CD31 (magenta), and PAX5 (white). Scale bar, 100 µm. **(G)** Representative GlycoScope multiplexed images of FL and FH showing DC-SIGNL (cyan), CD21 (magenta) and CD21 (yellow). scale bar: 100µm. **(H)** Violin plots showing the expression of DC-SIGNL acquired from GlycoScope in tonsil, FH and FL tissues (Krus-kal-Wallis rank sum test followed by pairwise Wilcoxon test with Benjamini-Hochberg adjustments). **(I)** Histogram showing the expression distribution of DC-SIGNL in B cell. **(J)** Violin plots showing the expression of BCL2 protein in DC-SIGNL^High^ and DC-SIGNL^Low^ B cells (Pairwise Wilcoxon test). **(K)** Line plot showing the mean expression of DC-SIGNL in follicle B cell at 0-150 µm from the anchored FDC. ***P<0.001; ****P<0.0001

We first evaluated the specificity of GlycoScope for glycan detection and compatibility with multiplexed imaging using a 6-plex panel of oligo-conjugated lectins (Siglec-7, -9, -15, Galectin-8, DC-SIGN, and MBL). We stained serial formalin-fixed paraffin-embedded (FFPE) human tonsil tissue sections with the lectin panel, with or without preincubation with their respective binding inhibitor. Each oligo-conjugated lectin exhibited a distinct membrane-staining pattern that was markedly diminished upon binding inhibitor treatment (**Fig. S5A**), suggesting that multiplexed imaging of these lectins enables specific glycan detection with minimal background noise and cross-reactivity.

To simultaneously detect glycans and proteins, we sequentially stained FFPE human tonsil tissues with our human lectins together with a 24-plex panel of oligo-conjugated antibodies encompassing various immune cell phenotypic markers (e.g., CD20, CD3, CD4) and then applied multiplexed imaging. Because antibodies are glycosylated, we compared the staining patterns and signal intensities across serial tissue sections for 1) lectins and antibodies together, 2) antibody only, or 3) lectin only. The glycan and protein staining patterns observed with GlycoScope mirrored those in the lectin-only or antibody-only conditions. We did not detect antibody signals in the lectin-only or lectin signals in the antibody-only conditions. These data indicate that the oligo-conjugated lectins do not cross-react with antibody glycans and that antibodies do not exhibit off-target binding to the oligo-conjugated lectins (**Fig. 4B and Fig. S5B**). We next evaluated the signal intensities of GlycoScope and observed no statistically significant differences for most glycan and protein markers compared to single-component (lectin or antibody only) conditions, respectively (**Fig. 4C, left**). A coefficient of variation (CV) score analysis, which quantifies the variability in staining intensities across groups, demonstrated strong concordance of GlycoScope staining as it was consistent with the antibody-only and lectin-only conditions (**Fig. 4C, right**). Thus, GlycoScope enables robust single-cell co-detection of glycans and proteins in FFPE tissue without compromising signal integrity from either individual modality.

### GlycoScope confirms mannosylated-BCR biology in follicular lymphoma

We next applied GlycoScope to interrogate glycan and immune organization in follicular lymphoma (FL), a lymph node-resident B cell malignancy that typically follows an indolent course but can undergo histologic transformation to transformed FL (tFL) or diffuse large B cell lymphoma (DLBCL) – events associated with markedly poorer prognosis. FL progression is driven in part by BCL2 overexpression and recurrent mutations that promote aberrant mannosylation of surface immunoglobulins or BCRs^36,37^. Mannosylated BCRs are proposed to engage soluble and membrane-bound mannose-binding lectins, including MBL and DC-SIGN, expressed on surrounding follicular dendritic cells (FDC) and macrophages, thereby sustaining BCR signaling and supporting malignant B cell survival^36–38^. Despite the importance of these pathways, the spatial distribution and functional consequences of lectin-binding glycans across malignant B cell states within intact FL tissue remain poorly defined, motivating the use of complementary spatial and single-cell glycomics approaches.

To address this issue, GlycoScope was applied to FFPE tissue micro arrays (TMAs) comprising four tonsils, four follicular hyperplasia (FH) samples, and eight FL cores, as well as a second cohort including five FH and eleven FL samples (**Fig. 4D and Supplementary Table 1**). Tissues were profiled using a 32-plex panel or 42-plex for cohort 2, encompassing phenotypic, activation/exhaustion, and FL-specific markers, together with six human recombinant lectins (Siglec-7, -9, MBL, DC-SIGN, Galectin-8, Galectin-9) (**Fig. S5C, Supplementary Table 2**). This design enabled simultaneous mapping of immune cell identity, functional state, and glycan landscape within preserved tissue architecture.

Using protein markers, we annotated six spatially organized immune subsets, including B cells, T cells, myeloid cells, endothelial cells, lymphatic cells, and follicular dendritic cell (FDC)–enriched niches (**Fig. 4E** and **Fig. S6A–B**). These annotations revealed well-defined spatial segregation of immune and stromal populations across tissue types. Consistent with established FL pathology, FL cores exhibited pronounced remodeling of the lymphoid architecture, including expansion of malignant B cell regions and reduced T cell infiltration compared with reactive tissues (**Fig. S6C-D**). Together, these findings validate the spatial fidelity of GlycoScope and its ability to capture known disease-associated tissue remodeling.

Building on this spatial framework, multiplexed glycan profiling by GlycoScope enabled direct visualization of the spatial distribution of glycans of interest within defined cellular compartments. Siglec-7L and Siglec-9L were preferentially localized to CD31^+^ endothelial cells across disease states (**Fig. 4F and Fig. S6E**). Although overall signal intensity was reduced in FL compared with reactive tonsil and FH tissues (**Fig. S6F**), the endothelial-restricted spatial distribution was preserved, indicating conservation of endothelial-associated glycan patterning with disease-associated modulation of signal magnitude.

We next focused on DC-SIGNL binding, a functional glycan feature with established relevance to FL. DC-SIGNL signal was significantly elevated in FL compared with FH (**Fig. 4G-H**), indicating disease-associated enrichment of DC-SIGN–accessible ligands. Stratification of malignant B cells by lectin-accessible glycan state (**Fig. 4I**) revealed that DC-SIGNL^High^ B cells exhibited markedly elevated BCL2 expression (**Fig. 4J**). This concordance between high DC-SIGN binding and BCL2 expression reflects hallmark molecular features of FL and is consistent with the model in which mannosylated BCRs engage DC-SIGN to reinforce BCR-dependent survival signaling, providing *in situ*, spatially resolved support for this mechanism^37,38^. We further quantified DC-SIGNL levels in follicular B cells as a function of distance from FDC-enriched niches (**Fig. 4K)**, and in both FH and FL, DC-SIGNL binding declined with increasing distance from FDCs, revealing a distance-dependent spatial pattern within follicles. Notably, FL samples maintained higher DC-SIGNL binding across all distances examined (**Fig. 4K**). Together, these data link FDC proximity to DC-SIGNL accessibility and suggest that FDC-B cell interactions may help sustain DC-SIGN-mediated signaling within the FL microenvironment.

To further validate these observations, we analyzed matched donor samples by scGOAT-seq on three tonsils, three FH samples, and five FL cases (**Fig. S6G**). In total, 31,924 high-quality immune cells were obtained and 18 different cell subtypes based on marker gene expression were identified (**Fig. S6H-J**). Across tonsil, FH, and FL samples, DC-SIGNL binding was elevated on malignant B cells and significantly enriched in FL compared with reactive tissues (**Fig. S6K-L**), mirroring GlycoScope patterns. GSEA showed that DC-SIGNL^High^ cells exhibited coordinated enrichment of pathways linked to antigen engagement and adaptive immunity, including antigen processing and presentation, immune response activation, and humoral and adaptive immune programs. In contrast, TNFα–NFκB signaling was significantly upregulated in DC-SIGNL^Low^ cells (**Fig. S6M**). Together, these spatial and single-cell datasets define a coherent FL glyco-state characterized by elevated DC-SIGNL accessibility, coordinated activation of BCR-associated malignant programs, and suppression of inflammatory NFκB signaling. More broadly, these results demonstrate that functional glycan states can be spatially organized, transcriptionally coupled, and quantitatively measured in human tumors, establishing functional glycan accessibility as a biologically meaningful dimension of the tumor micro-environment.

### GlycoScope reveals spatially organized Galectin-8 binding patterns in FL neoplastic follicles associated with localized immune cell organization

The multiplexed aspect of GlycoScope allows simultaneous interrogation of multiple glycans, enabling the discovery of coordinated glycan alterations in FL tissue. In addition to the expected increase in DC-SIGN binding in FL (**Fig. 4**), we observed an elevation of Galectin-8 interactions (**Fig. S6F**), suggesting that the glycan-lectin axes is dramatically altered in disease. Analysis of the glycan distributions across annotated cell types revealed a selective increase in Galectin-8 binding to Pax5+ B cells (**Fig. 5A-B**). Notably, Galectin-8 interactions were markedly increased in the neoplastic follicular B cells in FL, compared to B cells from both the mantle zone and follicle center in FH and tonsil (**Fig. 5C**), suggesting a spatial feature not previously described in this disease, despite prior evidence that other galectins are perturbed in B cell malignancies^42,43^.

**Fig 5.**
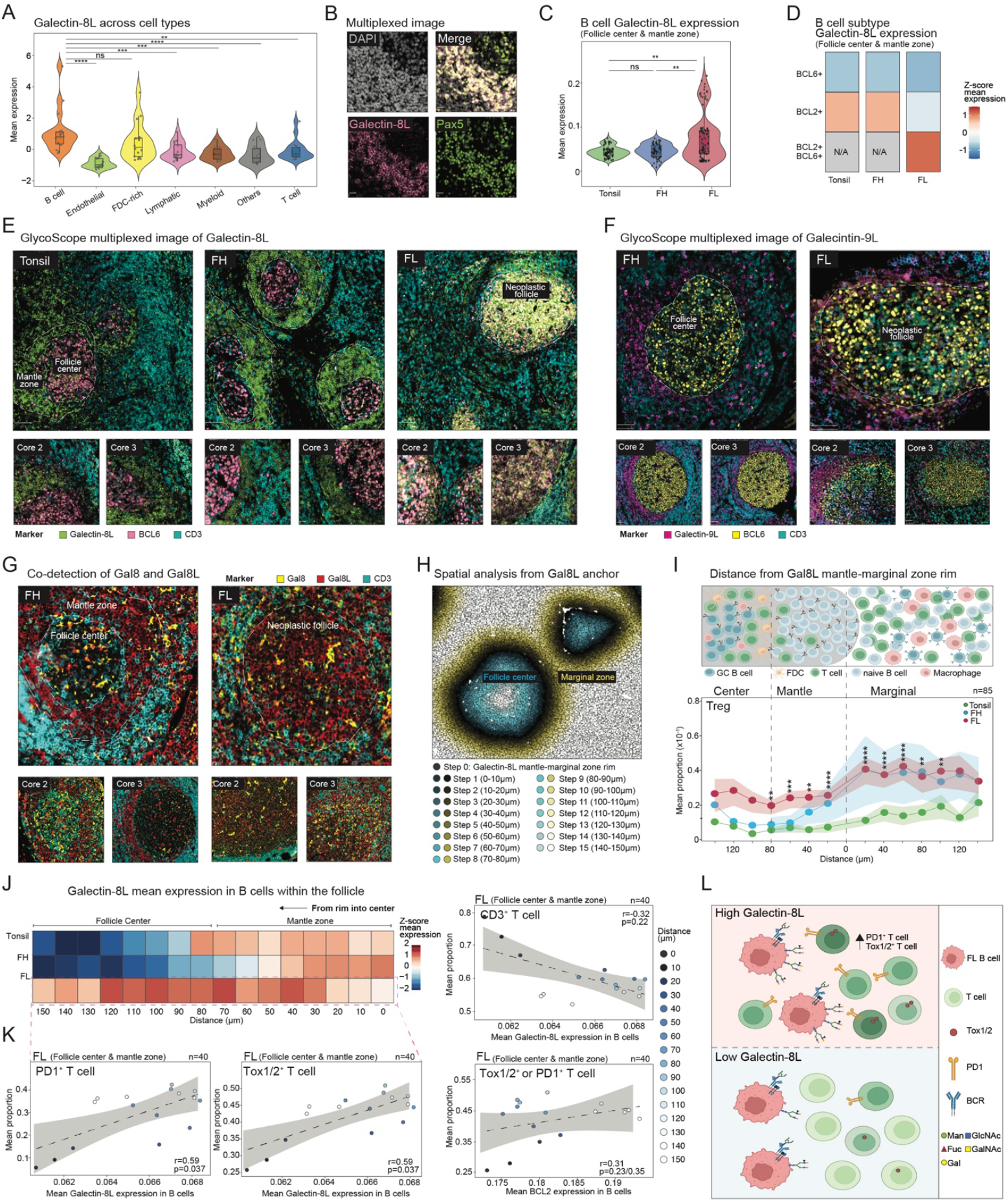
GlycoScope maps a distinct Galectin-8L spatial pattern in FL neoplastic follicles that associate with lectin proximity and distinct intrafollicular immune niches. **(A)** Violin plots showing the median expression of Galectin-8L across cell types (Kruskal-Wallis rank sum test followed by pairwise Wilcoxon test with Benjamini-Hochberg adjustments). **(B)** Representative GlycoScope multiplexed images with markers for Galectin-8L and B cells (Pax5) shown (scale bar: 20µm). **(C)** Violin plots showing the mean expression of Galectin-8L in B cells within the follicle across tonsil, FH, and FL tissues (Kruskal-Wallis rank sum test followed by pairwise Wilcoxon test with Benjamini-Hochberg adjustments). **(D)** Mean expression of Galectin-8L in BCL6+, BCL2+, and neoplastic BCL2+BCL6+ follicular B cells across tonsil, FH, and FL tissues. **(E)** Representative GlycoScope multiplexed images with markers for Galectin-8L (green), BCL6 (pink), and CD3 (cyan) shown (scale bar: 100µm). White dotted lines marked the regions for mantle zone, follicle center and neoplastic follicle. **(F)** Representative GlycoScope multiplexed images with markers for Galectin-9L (magenta), BCL6 (yellow), and CD3 (cyan) shown (scale bar: 100µm). White dotted lines marked the regions for follicle center and neoplastic follicle. **(G)** Representative GlycoScope multiplexed images with markers for Galectin-8L (magenta), Galectin-8 (yellow), and CD3 (cyan) shown (scale bar: 100µm). White dotted lines marked the regions for mantle zone, follicle center and neoplastic follicle. **(H)** Schematic illustration of stepwise extension analysis. Galectin-8L+ B cells at the mantle-marginal zone rim were anchored and expanded inward towards the follicle center (blue) and outward towards marginal zone (red), in 10 µm radial steps. **(I)** Mean proportions of Treg between 0-140µm towards the follicle center and marginal zone from the anchor rim are shown. The region in grey symbolizes the follicle. Mean proportions of FDC-enriched B cells, CD4+ T, CD8+ T, CD68+ macrophage and CD68+CD163+ macrophages are shown in **Fig. S6B-C** (Kruskal-Wallis rank sum test followed by pairwise Wilcoxon test with Benjamini-Hochberg adjustments). **(J)** Mean expression of Galectin-8L in follicular B cells at each step across tonsil, FH, and FL tissues. **(K)** Correlation scatter plot of the proportion of PD1+ (left) and Tox1/2+ (right) T cells with Galectin-8L expression in B cells at each step (distance towards follicle center from anchor rim) (Spearman’s correlation; p-value adjusted with Benjamini-Hochberg method). **(L)** Top: Correlation scatter plot of the proportion of CD3T cells with Galectin-8L expression in B cells at each step (distance towards follicle center from anchor rim). Bottom: Correlation scatter plot of the proportion of Tox1/2+ or PD1+ T cells with BCL2 expression in B cells at each step (distance towards follicle center from anchor rim) (Spearman’s correlation; p-value adjusted with Benjamini-Hochberg method). **(M)** Schematic illustration of summary findings. *P<0.05; **P<0.01; ***P<0.001; ****P<0.0001; ns P>0.05.

To further delineate the spatial identity of these Galectin-8L-interacting B cells, we integrated Galectin-8L complexation with canonical follicular structural markers BCL2 and BCL6. In tonsil and FH tissues, Galectin-8L interactions were largely confined to BCL2+ mantle-zone B cells, whereas in FL, they were enriched in neoplastic B cells co-expressing BCL2 and BCL6 (**Fig. 5D**). This organization was readily apparent in multiplexed images, where Galectin-8L-binding B cells were restricted to the mantle zone in tonsil and FH, but extended across both the follicular center and mantle regions in FL (**Fig. 5E** and **Fig. S7A**). This pattern was specific to Galectin-8L as Galectin-9L interactions were largely restricted from the follicle center and distributed similarly in FH and FL (**Fig. 5F**). Furthermore, co-detection of Galectin-8 accessible ligands and endogenous Galectin-8 lectin revealed that Galectin-8 lectin was uniformly distributed across follicles in both FL and FH (**Fig. 5G**). Thus, the distinct FL-specific distribution of accessible Galectin-8L, rather than altered lectin abundance, is indicative of the changes in disease.

Motivated by this FL-specific spatial patterning, we next examined whether Galectin-8L binding could distinguish follicular architecture. We thus performed stepwise neighborhood analysis from the Galectin-8L^High^ mantle-marginal zone rims, extending inward towards the follicular center and outward toward the marginal zone in 10 µm increments up to 150 µm (**Fig. 5H**)^44,45^. Using follicular dendritic cell (FDC)–enriched regions^46^ as internal landmarks, we analyzed the follicular center, mantle zone, and marginal zone (**Fig. S7B**). We defined the follicular center as the region located 80-150 µm inward from the Galectin-8L^High^ mantle zone rim, the mantle zone as spanning 0-80 µm, and the marginal zone as extending outward of the rim. This approach revealed disease-specific spatial distributions of immune populations. In FL, T regulatory cells (Tregs) were consistently elevated throughout the mantle zone compared to FH and tonsil. Strikingly, a sharp increase in Treg abundance was observed at the transition from mantle to marginal zones in FL and FH, followed by a plateau across marginal steps, whereas tonsil tissues maintained consistently lower levels throughout (**Fig. 5I**). In contrast, CD8+ T cells were enriched in FH and tonsil at multiple mantle and marginal positions but were markedly reduced in FL (**Fig. S7C**, left, top). CD68^+^ macrophages exhibited a progressive increase across the marginal zone in FL and FH (**Fig. S7C**, left, bottom), whereas CD4+ T cells (**Fig. S7C**, right, top) and CD68+CD163+ macrophages (**Fig. S7C**, right, bottom) generally showed similar trends of changes across regions and disease states.

Within FL follicles, Galectin-8 binding was not uniform; it displayed a spatial gradient, with elevated levels observed between 30–120 µm and lower expression at the inner and outer follicular boundaries (**Fig. 5J**). This heterogeneity prompted us to explore whether accessible Galectin-8L abundance correlated with the localized distribution of immune cells. Intriguingly, we observed that Galectin-8L levels in follicular B cells were positively correlated with the local abundance of PD1+ or Tox1/2+ T cells (**Fig. 5K**), indicating spatial clustering of these specific T cell populations near Galectin-8L^High^ regions within the follicle. This association was not observed for other checkpoint markers such as TIM3 and LAG3 (**Fig. S7D**), which showed negative correlations, nor for total CD3^+^ T cells (**Fig. 5K**, right), suggesting that the clustering is not due to general T cell infiltration. Furthermore, this spatial relationship is distinctly Galectin-8L-dominant, as BCL2 expression, despite its broad presence in neoplastic B cells in FL, showed no significant correlation with these T cell subsets (**Fig. 5K**, right). Together, these results show that Galectin-8L binding is spatially reorganized in FL and defines intrafollicular regions with distinct immune composition. By using Galectin-8L binding as a spatial landmark, GlycoScope resolves novel glycan-defined tissue architecture and reveals immune neighborhoods not captured by protein markers alone, establishing functional glycan accessibility as an additional dimension of tumor microenvironment organization. (**Fig. 5L)**.

## Discussion

Functional glycans shape immune recognition, cell– cell communication, and tumor–immune interactions, yet have remained largely inaccessible to single-cell and spatial profiling. Existing approaches capture glycan abundance or biosynthetic potential but rarely report how glycans are interpreted by endogenous receptors in physiological contexts. Here, we engineered a panel of recombinant human lectins compatible with multimodal single-cell and spatial platforms, enabling direct measurement of lectin-accessible glycan states using the same receptor families that engage these structures in vivo. Together, scGOAT-seq and GlycoScope enable high-throughput, biologically grounded glycan readouts that integrate with transcriptomics and spatial architecture, revealing glycan-defined cell states and tissue programs that are not resolved by transcriptional or protein markers alone.

Perturbation of sialic acid biosynthesis in primary naïve CD8+ T cells highlights a conceptual advantage of functional glycan profiling with scGOAT-seq. Disruption of GNE, a central enzymatic node in Neu5Ac biosynthesis, led to a marked reduction in Siglec-7 ligand accessibility, accompanied by coordinated changes in activation-associated transcriptional programs. These findings suggest that alterations in cell-surface sialylation are detected by endogenous lectins and can be linked to immune activation states, even in otherwise minimally stimulated naïve lymphocytes. Rather than acting solely as passive structural features, sialylated glycans appear to be coupled to activation-associated T cell states. In this context, scGOAT-seq provides a means to connect perturbations of glycan biosynthesis to immune state readouts at single-cell resolution, complementing existing transcriptomic and genomic approaches. More broadly, these data support the view that glycan regulation constitutes an additional layer of immune control that intersects with canonical immune signaling programs.

Across PBMC perturbations, endogenous lectin binding revealed stimulus-specific remodeling of cell-surface glycan accessibility that was not apparent from transcriptional programs alone. Siglec-9 and Siglec-15 binding stratified activated and effector-like CD4+ T cell states, whereas enhanced Siglec-7 interactions marked an IL-2–independent activation program most strongly induced by LPS. Notably, lectins within the Siglec family respond divergently to inflammatory cues: Siglec-7 binding increased following innate stimulation but decreased with TCR activation, while Siglec-9 and Siglec-15 interactions increased selectively with TCR stimulation and associated with cytokine- and IL-2–responsive effector programs.

These patterns are consistent with cell-type–restricted expression of Siglec family members, but further reveal that human lectins with closely related glycan specificities exhibit distinct and non-redundant cell-binding behaviors within the same immune populations. Despite recognizing overlapping classes of sialylated glycans, Siglec-7, Siglec-9, and Siglec-15 displayed stimulus-dependent and cell-type–specific binding, indicating that glycan remodeling is interpreted through distinct lectin interactions deployed by each immune population rather than reflecting glycosylation gene expression alone. These findings extend prior mechanistic studies of Siglec-7 and Siglec-9 signaling in human T cells^50,51^ and NK cells^32^ by placing them in a context-dependent glycan framework, in which distinct cues drive non-overlapping lectin-accessible glycan states. Notably, these stimulus-specific patterns were not apparent from the expression of glycosylation pathway genes, underscoring that endogenous lectins report functional glycan remodeling not captured by transcript abundance alone.

Flow cytometry confirmed the scGOAT-Seq findings that Siglec-ligand abundance correlated with cytotoxic effector states, supporting the biological relevance of these glycan-defined programs. These results suggest that cell-surface lectins can be used to enrich and sort T cell fractions with higher cytotoxicity, or to inhibit them. More generally, these results indicate that lectin-based readouts resolve an additional layer of immune regulation and imply that Siglec-directed therapeutic strategies may differentially impact T cell activation programs depending on the inflammatory milieu and the glycan states it induces.

Perturbation of Siglec–sialic acid recognition further suggested that Siglec engagement is coupled to broader remodeling of lectin-accessible glycan states during inflammation. Blocking Siglec-7 and Siglec-9 during LPS stimulation abolished the expected increase in Siglec-7L, reduced Siglec-9L, and altered accessibility of multiple lectin family members, including those from C-type lectins, galectins, and Siglec-15. These glycan shifts coincided with marked transcriptional rewiring in monocytes and NK cells, including the induction of NF-κB signaling, unfolded protein response programs, metabolic regulators, and antigen-presentation machinery. These patterns are consistent with reports that disrupting Siglec-7/9–sialic acid engagement alters inhibitory signaling and inflammatory reprogramming in myeloid and lymphoid cells^23,32,52^. Network-based inference further indicated that the effects of Siglec blockade extended beyond directly targeted cell types: whereas LPS stimulation alone produced a predominantly monocyte-centered cytokine network, Siglec blockade shifted inferred communication toward interferon-, cytotoxic-, and co-stimulatory programs involving NK and T cells. CD8+ T cells, despite lacking direct antibody engagement, exhibited secondary induction of interferon-stimulated genes and TNF-associated pathways, illustrating how perturbation of a single glycan–lectin axis can reshape distal immune states through altered cell–cell communication. Together, these findings support a model in which lectin–glycan interactions contribute to the organization of immune activation states.

Integrated spatial and single-cell glycomics further revealed glycan-defined malignant B cell states in FL. ScGOAT-seq and GlycoScope converged on two malignant FL populations distinguished by the engagement of mannose-binding lectins. DC-SIGNL^High^ malignant B cells were enriched for BCR-associated and germinal center–linked transcriptional programs, directly linking prior biochemical models^36,38^ of mannosylated BCR–DC-SIGN interactions to cell-state resolved and spatially defined malignant populations in human tissue. In contrast, DC-SIGNL^Low^ cells upregulated TNFα–NFκB inflammatory programs with reduced lectin binding, suggesting alternative malignant states with distinct signaling contexts.

Beyond the mannosylated BCR axis, GlycoScope uncovered a disease-associated spatial glycan pattern centered on Galectin-8 binding. In reactive lymphoid tissues, Galectin-8L detection was largely confined to mantle-zone B cells, whereas in FL it was broadly distributed across neoplastic B cells throughout the follicle. This redistribution was not seen for Galectin-9, indicating that altered galectin accessibility is not uniform across the family. Increased galectin binding is often associated with reduced terminal sialylation and greater exposure of LacNAc/galactose residues^54^, although lectin accessibility can also be shaped by glycan density and presentation. Notably, Galectin-8 protein was relatively uniformly distributed across follicles, suggesting that disease-associated changes in ligand accessibility, rather than lectin abundance, underlie Galectin-8L engagement in FL. Together, these observations demonstrate that functional glycan accessibility is spatially structured in tissue and can reveal disease-associated microenvironments not captured by transcriptomic or proteomic profiling alone.

Leveraging Galectin-8L as a positional anchor, GlycoScope resolved intrafollicular immune neighborhoods with distinct cellular composition. Regions enriched for Galectin-8L-expressing malignant B cells preferentially co-localized with PD-1+ and TOX+ CD4+ T cell subsets. defining a glycan-associated microenvironment not captured by conventional protein markers such as BCL2 alone. Together with the DC-SIGNL-associated mannosylation linked to BCR-active malignant B cells, these findings support a model in which glycan accessibility constitutes a critical layer of tissue organization in the FL microenvironment, operating alongside genetic and protein-based programs. More broadly, these results highlight lectin– glycan axes beyond the Siglec–sialic acid recognition that organize immune states in human tissues. Rather than acting as canonical inhibitory receptors, galectins and other lectin families may demarcate spatially organized glycan features associated with distinct immune cell neighborhoods.

Together, scGOAT-seq and GlycoScope provide a modular strategy for measuring human lectin-accessible glycan states within single-cell and spatial contexts. By enabling multimodal readout of lectin-recognized glycans alongside transcriptional programs and tissue architecture, these platforms extend standard single-cell and spatial workflows with a functional glycan dimension. Lectin binding reports ligand accessibility and avidity under assay conditions rather than complete glycan structures, motivating future integration of these methods with orthogonal glycomic measurements. Beyond the systems examined here, the modular design of scGOAT-seq and GlycoScope allows extension to additional lectins, perturbations, and disease contexts. Looking ahead, the modular design of scGOAT-seq and GlycoScope should enable expansion to additional lectins, perturbations and disease contexts, supporting applications in biomarker discovery, therapeutic stratification and the development of glyco-immune-targeted interventions.

## Acknowledgements

We thank Craig Lassy and Michael Hair from Akoya for Phenocycler Fusion technical support, Marissa Vera for providing DC-SIGN cDNA for plasmid cloning, Amanda Dugan, Nuo Liu, Sarah Quinn, Peter Winter, and Srivatsan Raghavan for helpful feedback and discussions.

This article reflects the views of the authors and should not be construed as representing the views or policies of the institutions that provided funding.

This paper was typeset with the bioRxiv word template by @Chrelli: www.github.com/chrelli/bioRxiv-word-template

## Author contributions

Conceptualization: AB, COC, SPTY, SJ, AKS, LLK

Experiment: AB, COC, SPTY, SA, SS

Analysis: AB, COC, SPTY

Contribution of reagents, tools, or technical expertise: input from all authors

Writing: AB, COC, SPTY, SJ, AKS, LLK

Supervision and funding: SJ, AKS, LLK

AB, COC, and SPTY reserve the right to list their names first in their CV.

## Competing interests

SJ is a co-founder of Elucidate Bio Inc., has received speaking honoraria from Cell Signaling Technology, and has received research support from Roche unrelated to this work. LLK reports compensation for scientific advisory board membership from Exo Therapeutics. AKS reports compensation for consulting and/or scientific advisory board membership from Honeycomb Biotechnologies, Cellarity, Ochre Bio, Relation Therapeutics, Parabalis Medicines, Passkey Therapeutics, Quotient Therapeutics, Danaher, IntrECate Biotherapeutics, Bio-Rad Laboratories, and Dahlia Biosciences unrelated to this work. The other authors declare no competing interests.

## Data and materials availability

All sequencing data will be deposited in GEO, imaging datasets in Zenodo, and analysis code in public GitHub repositories, and will be made available upon publication. For peer review, all data and code are currently hosted in a shared Dropbox folder.

## Methods and materials

### Human tissue acquisition and patient consent

#### GlycoScope

All formalin-fixed paraffin-embedded (FFPE) tissues used in this study were sectioned at 5 µm thickness on SuperFrost glass slides (VWR, 48311-703). The tonsil tissue block as part of GlycoScope technical benchmarking was commercially acquired (AMSbio, 194107) and sectioned by L.P. with 4 immediately adjacent sections each on the same slide.

#### FL/FH biological studies

The FL/FH TMA and their corresponding single-cell tissue suspensions for scGOAT-seq and GlycoScope studies were generously provided by W.R.B. from the University of Rochester Medical Center (IRB#STUDY00000159 - ULAB03012). To compare tonsil, FH, and FL as part of the GlycoScope analysis, 16 patient samples (4 tonsil, 4 FH, 8 FL) were sectioned from a tissue microarray (TMA) comprising a single 2.0 mm diameter core from each individual patient. In parallel, single-cell tissue suspensions from 10 patients (5 FH, 5 FL) were obtained for scGOAT-seq analysis. Both TMA and single-cell tissue suspensions were prepared by P.R. and W.R.B. (IRB#STUDY00000159 - ULAB03012). Detailed deidentified information for the FH and FL patients is available in **Supplementary Table 1** All patients were treatment-naive at the time of biopsy.

### Cloning and site-directed mutagenesis

Recombinant human Galectin-9 (Gal-9 mC10-HPPY)^63^ and human galectin-8 (Gene ID: 3964) were synthesized and cloned into a pET24a plasmid including an N-terminal AviTag™ and a C-terminal His6 tag by Twist Bioscience (Twist Bioscience).

Human MBL (*MBL2*) coding sequence (Uniprot ID P11226) was obtained from NCBI (gene ID: 4153) and synthesized as a DNA gene fragment (gblock) including an N-terminal AviTag™ and a His_6_ tag (Integrated DNA Technologies). DC-SIGN (CD209, UniProt ID Q9NNX6) was PCR out of Raji/DC-SIGN cells using the following primers (primers reverse 5′-GGGTTCCCTCCCACAAAACTCCTACGCAG-GAGGGGGGTT -3′ and forward 5′-GTCACACCCACAATTTGAAAAACAGCTTAAAGCTG-CAGTG -3′). The DNA fragments were Gibson ligated into a linearized pcDNA4 plasmid (primers reverse 5′-GGTGAA-GCTTAAGTTTAAACG-3′ and forward 5′-TGAG-GATCCACTAGTCCAGTG-3′). To insert an N-terminal AviTag™ and His_6_ tag, a gblock was obtained including 5′-TGTAATCTTGTCTGAGTCTCAGC -3′ and forward 5′-AGCTCTCAGAGAAATCCAAGC 3′ homology arms and cloned by gibson assembly into the linearized pcDNA4 DC-SIGN plasmid. Correct insertion was verified by Sanger sequencing (Quintara Biosciences).

### Protein expression and purification

Galectin-8 and Galectin-9 were expressed in D3 *Escherichia coli* from a pET-24a vector. Liter-scale cultures were grown to the mid-log phase (OD_600_=0.3-0.5) and induced with 0.4 mM isopropyl β-thiogalactoside (IPTG). Cultures were then grown for 16 hours at 18 °C and pelleted by centrifugation (20 min at 3000xg). After storage at -80 °C, pellets were resuspended in Ni-NTA Loading Buffer (20 mM sodium phosphate, 500 mM sodium chloride, 20 mM imidazole, pH 7.4) and lysed by French press (two cycles, maximum pressure 1500 bar) or B-PER II Bacterial Protein Extraction Reagent (Thermo Fisher 78260). Galectin-9 was supplemented with cOmplete™ protease inhibitor cocktail (Roche 04693116001) ^63^. Cell debris was removed by centrifugation (30 min at 33,000xg). DC-SIGN and MBL were each expressed transiently into Expi293F cells (Thermo Fisher Scientific) using ExpiFectamine293 Transfection Kit according to the manufacturer’s instructions (Thermo Fisher Scientific A14524). After 3 days, the conditioned expression medium was harvested by centrifugation and passed through a 0.2 mm PES filter.

Filtered supernatants were then loaded onto a Ni-NTA affinity column (BioRad, Hercules, CA), washed in Ni-NTA Loading Buffer, and then eluted on a gradient to Ni-NTA Elution Buffer (20 mM sodium phosphate, 500 mM sodium chloride, 500 mM imidazole, pH 7.4). Fractions were analyzed on a stain-free tris-glycine gel. Following purification, proteins were dialyzed in PBS with SnakeSkin 7,000 MWCO dialysis tubing (Thermo Scientific 68700) overnight at 4 °C to remove excess imidazole. Protein concentrations were determined by absorbance at 280 nm. Extinction coefficients and molecular weights were calculated using the ProtParam tool (web.expasy.org/protparam). Proteins were verified to be stably folded via differential scanning fluorimetry or circular dichroism spectroscopy before experiment use.

### Protein biotinylation

All lectins were biotinylated using an Enzymatic Protein Biotinylation Kit (Sigma-Aldrich CS0008) following the manufacturer’s instructions using a 1:100 BirA/Protein ratio (w/w) at 30 ºC for 1 hour. Excess biotin was removed using a Zeba 1.5 ml 7 K MWCO Spin Desalting Column (Thermo Fisher Scientific 89882) following the manufacturer’s protocol. Biotinylation was confirmed by a streptavidin gel-shift assay. Biotinylated human Siglec-7 (SG7-H82E7), Siglec-9 (SI9-H82E9), and Siglec-15 (SG5-H82E9) were obtained from Acro Biosystems.

### Cell lines

H6c7 (Kerafast) cells were cultured in KGM™ Gold Keratinocyte Growth Medium BulletKit™ (LONZA 192060). AsPC-1 (ATCC), A549 (ATCC) and Toledo (ATCC) cells were cultured in RPMI ATCC modified media (Gibco A1049101), both containing 10% fetal bovine serum (FBS, Gibco 10082147) and 1% Penicillin-Streptomycin (P/S, Gibco 15140122) at 5% CO2 and 37 °C.

### Isolation of PBMCs

PBMCs were isolated from Leukocyte Reduction System cones (STEMCELL Technologies 200-0093) which were processed within 48 h of the initial blood draw (**Supplementary Table 1**). After initial wash, leukocytes were layered gradually onto 15 ml of Lymphoprep™ (STEMCELL Technologies 07801). Samples were centrifuged at 1200xg in a table-top swinging bucket centrifuge for 10 min. PBMCs were carefully isolated from the Lymphoprep™ interface and added to 40 ml of PBS/FBS before being centrifuged for 8 min at 800xg. After the supernatant was discarded, a second wash was performed. PBMCs were frozen in Cryostor CS10 (BioLife Solutions 210373) overnight at −80 °C in a cryogenic freezing chamber. PBMCs were then transferred to liquid nitrogen storage for long-term storage.

### Immune Perturbations

PBMCs (n=3) were thawed and rested in Advanced RPMI supplemented with 10 % FBS, L-glutamine, HEPES and penicillin streptomycin for 2 hours at 37°C, after which they were seeded into 96 well plate with 200,000 cells in 200ul media. For LPS stimulation, 4ul of 1:100 LPS (Thermofisher Catalog# 00-4976-03) were added to each well and then cultured for 4 hours at 37°C. For antibody blocking experiments, cells were blocked with Fc blocking reagent, Human TruStain FcX (BioLegend Catalog# 422302) for 15 min followed by addition of human Siglec7/CD328 antibody (MAB1138-SP; R&D Systems) and human Siglec 9 antibody (MAB1139-SP; R&D Systems) each at a final concentration of 10ug/mL followed by LPS stimulation described as above. For TCR stimulation, PBMCs were rested for 2 hours after thawing and then stimulated with Dynabeads CD3/CD28 for human T cell activation (Thermofisher Catalog# 1131D) at a ratio of 1:1, followed by incubation at 37°C for 24 hours. After incubation, beads were separated using magnetic separation and cells were counted and labelled with oligo-lectin constructs for scGOATseq in cell staining buffer supplemented with 1mM calcium chloride.

### CRISPR-KO in Naïve CD8+ T cells

After thawing 100 M PBMCs for 3 independent biological donors, naïve CD8+ T cells were isolated using naïve CD8+ T cell isolation kit (STEMCELL 17968) as per manufacturer’s instructions. Cells were rested in media at 37°C for 4 hours. Guide RNAs for GNE gene (Guide1: 5’-mG*mG*mA*rCrCrArUrCrGrCrArUr-3’; Guide2: 5’-mC*mG*mA*rArUrCrCrUrUrCrArCrArUrUr-3’) and scrambled control were obtained from IDT and mixed with nuclease at 2.5:1 guide: nuclease molar ratio with a concentration of 0.2 mg/mL. CD8+ T cells with guides (1:1 ratio for Guide1 and Guide2) and scrambled control RNPs were then subjected to mechanoporation based gene delivery using the Portal Biosciences T cell boosting protocol as per manufacturer’s instructions. Cells after boosting were then counted and labelled with oligo-lectin constructs for scGOATseq downstream processing.

### Direct labeling of lectins and streptavidin

Streptavidin (Invitrogen 434301) was functionalized using DBCO-PEG4-NHS Ester (Vector Labs CCT-A134) dissolved in dimethyl sulfoxide at 1:10 in PBS pH 7.4. The reaction was incubated for 30 min at room temperature with mixing. To stop the reaction, 50 mM Tris-HCl, pH 8.0 was added. Non-reactive DBCO was removed by desalting using a Zeba 1.5 ml 7 K MWCO Spin Desalting Column (Thermo Fisher Scientific 89882) following the manufacturer’s protocol. DBCO labeling was determined using the following formula: A309 × εStreptavidin/((A280 − CF × A309) × εDBCO), where CF = 1.089, εStreptavidin = 41326 cm−1 M−1 and εDBCO = 12000 cm−1 M−1. 5’ azide-modified oligos (Integrated DNA Technologies) in **Supplementary Table 2** were incubated with DBCO-PEG4-Streptavidin 10:1 in PBS for 18 h with mixing at 4 ºC. Samples were purified using Amicon Ultra 3 kDa (Millipore Sigma UFC5003) to remove excess oligos. Single-strand DNA concentration was quantified using Qubit™ ssDNA Assay Kit (Thermo Fisher Scientific Q10212).

### Flow cytometry

All centrifugation steps were performed at 400 relative centrifugal force (RCF). Adherent cells were harvested using Trypsin-EDTA (Gibco 25200072). Collected cells were washed with DPBS (Thermo Fisher Scientific 14190250) and then resuspended in sterile filtered staining buffer made of 10% FBS with 1 mM CaCl_2_ in DPBS (Gibco 14040133). Staining with individual lectins was performed on 10e^5^ cells incubated with 1 µg/ml (DC-SIGN) or 2 µg/ml of human lectins and 0.5 µg/ml of Streptavidin Alexa Fluor™ 647 or 488 (Thermo Fisher Scientific S21374) for 40 min at 4 °C. Cells were subsequently centrifuged and washed with staining buffer twice. Cells were analyzed on an Attune Nxt flow cytometer (Thermo Fisher Scientific). Data was analyzed using FlowJo LLC software.

To quantify similarity between single-cell fluorescence distributions, we computed the Bhattacharyya coefficient (BC) for each pairwise comparison. Single-cell events were treated as independent observations of a one-dimensional fluorescence distribution for the specified channel. All gating, compensation, and quality filtering were performed upstream using FlowJo.

Probability density estimates for each sample were generated using Gaussian kernel density estimation (R function `density`, `bw=`“`nrd0`”, 512 points). Densities were evaluated on a shared grid spanning the intersection of the two distributions. Each probability vector was normalized to integrate to one. BC was calculated as:

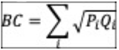

Where 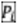 and 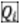 represent the normalized densities of the two samples on the common grid. BC values range from **0** (no overlap) to **1** (identical distributions).

### Multicolor Spectral Flow Cytometry of TCR stimulated T cells

PBMCs form 6 independent biological donors were thawed in media and T cell isolation was performed using the Stem-Cell EasySep Human T cell isolation kit (StemCell Catalog #17951) as per manufacturer’s protocol. Following T cell isolation, cells were counted and 1e^6^ cells were stimulated with Dynabeads CD3/CD28 for human T cell activation (Thermofisher Catalog# 1131D) at a ratio of 1:1 in 1ml media in 24 well plates for overnight at 37°C. After that, cells were activated using Cell Activation cocktail (with Brefeldin A) (Biolegend #423304) at a dilution of 1:500 for 4 hours. Cells were then separated from beads using magnetic separation and 200,000 cells were seeded per well in 96 well plate for staining. Cells were stained in staining buffer of 200ul with 15uM lectins and 10uM Alexa Fluorophore 488 conjugated streptavidin (Thermofisher Catalog# S32354) for 40 minutes at 4 °C. After washing, cells were stained with viability dye (Zombie R685 Viability kit, Biolegend Catalog #423119) and then washed. Cells were stained with cell surface protein antibodies at a dilution of 1:100 by volume (Spark PLUS UV 395 anti-human CD4 antibody, Biolegend #344628; Brilliant Violet 605 anti-human CD3 antibody, Biolegend #300460; PerCP anti-human CD69 antibody, Biolegend #310928; Brilliant Violet 421 anti-human/mouse/rat CD278 (ICOS) antibody, Biolegend #) for 30 min at 4 °C. Cells were then washed and permeabilized for 30 min at 4 °C, permeabilized and stained for intracellular cytokine antibodies at a dilution of 1:100 for 30 min at 4 °C (Brilliant Violet 785 anti-human IFNγ antibody, Biolegend #502542; PE anti-human Perforin antibody, Biolegend #353304, RB613 mouse anti-human Granzyme B antibody, BD #571117) followed by washing. Cells were then profiled for flow cytometry using Cytek Aurora spectral flow cytometer with single color controls and then analyzed using FlowJo software.

### Liquid Ligand Array (LiGA)

Streptavidin-coated, pre-blocked 96-well plates (Ther-mofisher, Cat#15501) were used to immobilize biotinylated lectins and capture binding phage from the LiGA library^18^. Biotinylated lectins were diluted to 25 µg/mL in 1× PBS, and 100 µL was added per well. Each lectin was analyzed in triplicate. Plates were sealed and incubated overnight at 4 °C. The coating solution was removed by inversion and blotting on absorbent paper. To block residual streptavidin sites, 100 µL of 10 mM biotin was added and incubated for 1 h. Wells were emptied and washed three times with 200 µL HBS (20 mM HEPES, 150 mM NaCl, 2 mM CaCl_2_, pH 7.4).

LiGA was diluted to 1×10^9^ PFU/mL in HBS, and 100 µL (≈1×10^8^ PFU) was added per well and incubated for 2 h. After incubation, wells were washed twice with HBS + 0.1% Tween-20 and once with HBS. Bound phages were eluted with 100 µL of 10 mM HCl. At exactly 9 min, eluates were transferred into tubes containing 50 µL Phusion 5× HF buffer for neutralization. A 2 µL aliquot of the neutralized eluate was used for qPCR to quantify enrichment. All steps were performed at room temperature unless noted. PCR protocol and Illumina sequencing and data processing was performed as previously described^18^.

### Cell mixing labeling

ASPC1and H6c7 cell lines were respectively stained following manufacturer’s instructions with CellTrace™ Far Red (Thermo Fisher Scientific C34564),and Violet (C34571) in 1 ml of DPBS for 20 min at 37 °C. Afterwards, cells were washed with DPBS (Thermo Fisher Scientific 14190250) and then resuspended in sterile filtered staining buffer made of 10% FBS with 1 mM CaCl_2_ in DPBS 100,000 cells/ml. Cells were mixed 1:1:1 before staining with individual Siglecs (2 µg/ml) and 0.5 µg/ml of Streptavidin Alexa Fluor™ 488 (Thermo Fisher Scientific S21374) for 40 min at 4 °C. Each cell mixing experiment was performed in triplicate.

### Sialidase treatment

Cells were treated with 50 mU/ml of the neuraminidase from *Arthrobacter ureafaciens* (Roche 10269611001) for 30 min at 37 °C.

### scGOAT-seq sample processing using Seq-Well custom streptavidin/TotalSeq A oligos

LDTs were generated by incubating oligo-conjugated streptavidin made in-house or TotalSeq A™ streptavidin PE probes (**Supplementary Table 2**) following manufacturer’s instructions with recombinant biotinylated human lectins on ice. After 15 min, biotin was added to block free binding sites and incubated for 15 min.

Pooled LDTs were added to 10e^5^ cells and incubated at 4 °C for 40 min in a staining buffer or 10% FBS (Thermo Fisher Scientific), 1 mM CaCl_2_ in RPMI media (Thermo Fisher Scientific; RP10). Afterward, the cells were centrifuged, washed with staining solution, and diluted to 10e^5^/ml. Massively parallel picowell based scRNA-seq was performed via the Seq-Well S^3^ platform as previously described^64,65^. Briefly, approximately 20,000 cells in 200 µl of staining buffer were loaded onto each PDMS based Seq-Well array, pre-loaded with barcoded RNA capture beads from ChemGenes. Arrays were rested for 10 min to allow gravity-based loading of cells into wells before being washed with PBS followed by non-supplemented media (RPMI) to remove any protein bound to the array surface and allow adequate membrane sealing. A plasma-treated semi-permeable membrane was added to each array and allowed to seal in an Agilent clamp at 37°C for 30 min. Arrays were then moved to lysis buffer (5 M guanidine thiocyanate, 10 mM EDTA, 0.1% BME, and 0.1% sarkosyl) for 20 min followed by hybridization buffer (2M NaCl, 8% v/v PEG8000) for 40 min. The arrays were then moved to wash buffer (2M NaCl, 3 mM MgCl2, 20 mM Tris-HCl, and 8% v/v PEG8000), and beads were removed from wells via pipette washing and manual scraping with glass slide. The removed beads then underwent reverse transcription at 52°C overnight using Maxima H Minus Reverse Transcriptase (ThermoFisher). Excess primers were then removed using Exonuclease I enzyme (New England Biolabs), denaturation with 0.1M NaOH was performed for 5 min followed by second strand synthesis using Klenow Fragment (3’5’ exo-) (New England Biolabs), and whole transcriptome amplification was achieved via PCR with KAPA HiFi Hot-Start ReadyMix (Roche), within spike-in primer for amplifying Seq-Well captured LDTs. cDNA and LDT libraries were separated during a 0.6X SPRI using AMPure XP beads (Beckman Coulter), where the beads captured the cDNA and the supernatant contained the LDTs. cDNA libraries were prepared using Nextera Dual 22 Index kits. The LDT libraries were further SPRI cleaned and then amplified using P5 and RPI P7 primers. The libraries were quantified and then loaded onto NextSeq 2000 P3 50 cycle kits and sequenced using the read structure: 21-8-0-50 for Read1-Index1-Index2-Read2. The resultant data was converted to FASTQ files using bcl2fastq and then aligned to the human genome (GRCh38) using drop-seq tools, on Terra maintained by the Broad Institute. Collapsing the aligned reads based on barcode and UMI sequences, digital gene expression matrices were obtained for each array, covering 10,000 cell barcodes.

### scGOAT-seq sample processing using 10X 5’ capture kit

Barcoded lectin panels were pre-conjugated with Biolegend TotalSeq-C Streptavidin probes for 15 min at 4 °C followed by quenching of free sites of streptavidin with free biotin for additional 15 min after which all barcoded lectin panels were pooled together. Cells were stained with barcoded lectin panels for 40 min at 4 °C, followed by washing. Cells from different donors were hashed together using Biolegend TotalSeq C hashing antibodies as per manufacturer’s instructions. Briefly, cells were first blocked with Fc-blocking reagent for 10 min, washed and then stained with hashing antibodies for 30 min, washed and pooled together. Cells were then counted and loaded onto 10X Chromium and down-stream protocols were performed as per manufacturer’s instructions for 10X Chromium 5’gene expression with cell surface antibody libraries. After cDNA and ADT/Hash libraries were prepared, they were quantified and loaded into Illumina P4 100 cycle NextSeq kits with a targeted molar ratio of 9:1 for sequencing. The resultant data was converted to FASTQ files using bcl2fastq and then aligned to the human genome (GRCh38) using 10X Cell Ranger pipelines, on Terra maintained by the Broad Institute.

### Data Analysis: scGOAT-seq

#### Data Processing and quality control (scGOAT-seq Seq-Well S^3^)

For individual samples, collapsed gene expression profiles from 10,000 cell barcodes were converted into Seurat V4 objects using the “CreateSeuratObject” function. Cells were filtered on the basis of the number of genes, number of features, and percentage of mitochondrial genes based on the distribution for each different experiment and sample type profiled. The percentage of mitochondrial gene expression was determined using the PercentageFeatureSet() function of Seurat with the pattern set to ^MT. For the cell-line mixture model comparison with flow cytometry, we filtered cells expressing fewer than 1,500 genes and more than 20,000 genes, as well as cells expressing more than 5% of their transcriptome as mitochondrial genes. For the PBMC and FL samples across donors, we filtered out cells with number of genes less than 300 and more than 4,000, number of UMIs less than 500 and more than 10,000, or expressing more than 25% of their transcriptome as mitochondrial genes. Data was normalized, scaled using default Seurat pipeline and the top variable features were identified using the FindVariableFeatures() function of Seurat with default parameters. Principal components analysis (PCA) was performed using RunPCA() with default parameters and an elbow plot using the ElbowPlot() function with ndims=40 to determine the dimensions for clustering and UMAP generation. UMAPs were generated using RunUMAP() with the PCA reduction. FindNeighbors() and FindClusters() functions were used for clustering, with a 0.8 resolution parameter. The LDTs were added as a separate assay to the Seurat objects, where each the counts for LDT per cell was normalized to the counts of the control streptavidin and then normalized using log-transformation. The FindMarkers() function was used to determine marker genes using default parameters for manual annotation of clusters into cell types. For the reference mapping, the FindTransferAnchors(), FindIntegrationAnchors() and the IntegrateData() functions were used to map clusters to the reference dataset atlas for annotation.

#### Data Processing (scGOAT-seq 10X 5’)

For the individual samples, collapsed gene expression profiles obtained from CellRanger were converted into Seurat V5 objects using the “CreateSeuratObject” function. For each particular sample, demultiplexing of the hash antibodies was performed using the Seurat HTODemux function with a positive quantile value of 0.90, following which doublets and empty droplets were filtered out for downstream analyses. The LDTs were added as a separate assay to the Seurat objects, where the counts for LDT per cell for each lectin were divided by the counts for the control streptavidin signal and then log-normalized. For gene expression data, data was log-normalized and scaled using default Seurat pipeline. Principal components analysis (PCA) was performed using RunPCA() with default parameters and an elbow plot using the ElbowPlot() function with ndims=35 to determine the dimensions for clustering and UMAP generation. UMAPs were generated using RunUMAP() with the PCA reduction. FindNeighbors() and FindClusters() functions were used for clustering, with a 0.5 resolution parameter. The LDTs were added as a separate assay to the Seurat objects, where each the counts for LDT per cell was normalized to the counts of the control streptavidin and then normalized using log-transformation. Cell type annotation was performed using reference annotation through the Azimuth pbmcref dataset on Seurat and manually checked by plotting reference celltype marker genes from Azimuth on predicted cell types.

#### Differential expression testing and gene set enrichment analyses

The cells were binned into lectin-binding high and lectin-binding low populations using manual cutoffs for each lectin based on their distribution. Pairwise differential gene expression was performed using the FindMarkers() function from Seurat using default parameters. The differentially expressed genes were ranked by log-fold change and the ranked list was used gene set enrichment analyses using the fgsea package using the Hallmark and/or manually curated reference pathways.

#### Cell-cell interaction analyses

Inferred cell-cell interaction analyses were performed using the NicheNet package for intercellular communication modelling across different conditions for PBMCs using default parameters and visualization of top 50 inferred interaction scores using circos plots.

#### Survival analyses

For the survival analyses with gene sets, a two‐step pipeline was implemented where, first, genemodule scores were computed using the Seurat AddModuleScore function with 100 “control” genes. Then, for survival analyses, module‐scores were normalized across patients, dichotomized by upper/lower quantiles (upper and lower 40th quantiles for DLBCL, quartiles for all other cases), and Kaplan–Meier estimation was performed with logrank testing for significance testing.

#### Bulk RNA-seq analyses

Publicly available bulk RNA-seq data from TCGA were converted into Seurat V4 objects using the “CreateSeuratObject” function and log-normalized. The gene(s) of interest were visualized using the VlnPlot() with a Wilcoxon rank-sum test performed using the ggsignif package in R.

### GlycoScope

#### Antibody panel selection, conjugation, and titration

Antibodies used in the tonsil and FH/FL studies were obtained from their respective commercial sources and were conjugated either in-house or through collaboration with Cell Signaling Technologies (CST). These include previously validated antibody clones^66–68^. For antibodies conjugated inhouse, the specificity of antibody candidates were first validated by immunohistochemistry (IHC) on FFPE lymphoid tissues to ensure robustness of staining. The selected antibody clones were then conjugated by maleimide-based chemistry with minor modifications from a previously published protocol^69^. Briefly, 50 or 100 µg of carrier-free antibody was concentrated using a PBS-T pre-wetted 50kDa filter (Sigma Millipore, UFC5050BK) and then incubated with 0.9 µM TCEP (Sigma, C4706-10G) for 10-30 min in a 37°C water bath to reduce the thiol groups for conjugation. Reduction was quenched by two washes with Buffer C (1 mM Tris pH 7.5, 1 mM Tris pH 7.0, 150 mM NaCl, 1 mM EDTA) supplemented with 0.02% NaN_3_. Maleimide oligos were resuspended in Buffer C supplemented with 250 mM NaCl. The reduced antibody was then incubated with 100 or 200 µg (for 50 or 100 µg of antibody, respectively) of maleimide oligos (Biomers, 5’-Maleimide) in a 37°C water bath for 2 hours. The resulting conjugated antibody was purified by washing five times with the 50kDa filter with high-salt PBS (1× DPBS, 0.9 M NaCl, 0.02% NaN_3_). The conjugated antibody was quantified in IgG mode at A280 using a NanoDrop (Thermo Scientific, ND-2000). The final concentration was adjusted by adding >30% v/v Candor Antibody Stabilizer (FisherScientific, NC0414486) supplemented with 0.2% NaN_3_, and the conjugated antibody was stored at 4°C. Each conjugated antibody was titrated and validated via immunofluorescence on FFPE lymphoid tissues. All final stains were reviewed with board certified pathologist W.R.B..

#### FFPE tissue rehydration, antigen retrieval and blocking

FFPE tissue slides were baked in an oven (VWR, 10055-006) at 70°C for 1 h and thoroughly deparaffinized by immersing in xylenes for 2× 5 min. The slides were then rehydrated by a series of graded solutions using a linear stainer (Leica Biosystems, ST4020), with each step proceeding for 3 min: 3× xylene, 2× 100% EtOH, 2× 95% EtOH, 1× 80% EtOH, 1× 70% EtOH, 3× UltraPure water (Invitrogen 10977-023), and finally left in UltraPure water (Invitrogen 10977-023).

Unless otherwise specified, all washes mentioned below were with gentle agitation. Antigen retrieval was performed at 97°C for 20 min with pH 9.0 Target Retrieval Solution (Agilent, S236784-2) using a PT Module (ThermoFisher, A80400012), after which the slides were cooled to room temperature on the benchtop and washed once in 1× PBS for 5 min. Tissue regions were circled with a hydrophobic barrier pen (Vector Laboratories, H-4000) and then washed in 1× TBS-T for 5 min. The slides were then equilibrated by washing in EDTA-free S2 Buffer (0.5× DPBS, 0.25% BSA, 0.02% NaN_3_, 250 mM NaCl, 61 mM Na_2_HPO_4_, 39 mM NaH_2_PO_4_) for 20 min.

Tissue slides were then subjected to biotin blocking with the Streptavidin/Biotin Blocking Kit (Vector Laboratories, SP-2002). Briefly, slides were first incubated with the streptavidin solution for 15 min at RT in a humidity chamber followed by 1x PBS washes for 5 min, twice. The slides were then incubated with the biotin solution for another 15 min at RT in a humidity chamber followed by the same washing steps. The slides were then blocked with Blocking Buffer that contains BBDG (5% normal donkey serum, 0.05% NaN_3_ in 1× TBS-T wash buffer (Sigma, 935B-09)) supplemented with 50 µg/ml mouse IgG (diluted from 1 mg/ml stock (Sigma, I5381-10mg) in S2), 50 µg/ml rat IgG (diluted from 1 mg/ml stock (Sigma, I4141-10mg) in S2), 500 µg/ml sheared salmon sperm DNA (ThermoFisher, AM9680), 50 nM oligo block (diluted from stock with 500 nM of each oligo in 1× TE pH 8.0 (Invitrogen, AM9849) and 1X Carbo-free™ Blocking Solution (diluted from 10X (Vector Laboratories, SP-5040-125)). Blocking was performed in a humidity chamber on ice during which was photobleached using Happy Lights (Verilux, VT22) for 1 h, with temperature kept below 40°C.

#### Antibody and lectin staining

Each conjugated antibody in the panel was first centrifuged at 12,500 rpm for 8 min at 4 °C to remove aggregates. They were then diluted in a single tube containing 300 µl Antibody Diluent (5% normal donkey serum, 0.05% NaN_3_ in 1x TBS-T) at a concentration based on prior titration. The antibody mixture was then filtered through a EDTA-free S2 Buffer pre-wetted 50kDa filter (Sigma Millipore, UFC5050BK) at 12,500 rpm for 8 min at 4°C. The filtered mixture was adjusted to the desired final volume by adding 25% v/v total volume with FFPE Block (50 µg/ml mouse IgG (Sigma, I5381-10mg), 50 µg/ml rat IgG (Sigma, I4141-10mg), 500 µg/ml sheared salmon sperm DNA (ThermoFisher, AM9680), 50 nM oligo block (diluted from stock with 500 nM of each oligo in 1× TE pH 8.0 (Invitrogen, AM9849) in S2) and topping up the remainder with Antibody Diluent. The antibody mixture was then filtered through a EDTA-free S2 Buffer pre-wetted 0.1 µm pore size filter (Millipore, UFC30VV00) at 12,500 rpm for 1 min at 4°C. Blocked/photobleached tissues were then incubated with the filtered antibody mixture at 4°C in a humidity chamber overnight.

For lectin staining, tissues were washed twice in GlycoBuffer for 5 min each at RT. The slides were then rinsed quickly with 1x PBS twice and washed once with 1x PBS for 2 min. The slides were fixed with 4% PFA (diluted from 16% stock (EMS Diasum, 15740-04) in 1x PBS (diluted from 10x stock) in a humidity chamber protected from light, at RT for 10 min. The slides were then rinsed quickly with 1x PBS twice and washed once with 1x PBS for 2 min. After that, the slides were equilibrated by washing twice in GlycoBuffer for 5 min each at RT.

Oligo-conjugated recombinant lectins were prepared by incubating recombinant biotinylated lectins with oligo-conjugated streptavidin in individual tubes at ratios optimized based on prior titration. The reactions were carried out in custom Antibody Diluent Blocking Solution, on ice and protected from light, for 15 min. To block free binding sites, biotin was added at one-third of the total reaction volume, followed by an additional 15-min incubation. Each of the oligo-conjugated recombinant lectins were pooled, applied to the antibody-stained tissue and incubated in a humidity chamber protected from light at 4°C overnight. For experiments that involved glycan manipulations as part of GlycoScope technical validation, 10 mM pH 8.0 EDTA (diluted from 0.5 M stock (Invitrogen, 15575020)), 20 mM lactose (diluted from stock 200 mM stock (Sigma Aldrich, 17814) and 100 mU/ml sialidase (diluted from 1 U/ml stock (Roche, 10269611001)) was added to the DC-SIGN and MBL, Galectin-8, and Siglec-7, Siglec-9 and Siglec-15 experiments, respectively. They were then applied to the tissue and incubated in a humidity chamber protected from light at 4°C overnight.

After incubation, tissues were washed twice in GlycoBuffer for 2 min each at RT. The slides were fixed with 1.6% PFA (diluted from 16% stock (EMS Diasum, 15740-04) in GlycoBuffer in a humidity chamber protected from light, at RT for 10 min. The slides were then rinsed quickly with GlycoBuffer twice and washed once with GlycoBuffer for 2 min. The slides were fixed with ice-cold methanol for 5 min on ice, followed by rinsing with GlycoBuffer twice and washed once with GlycoBuffer for 2 min. The slides were then fixed in 4 µg/µl of BS3 Final Fixative (diluted from 200 µg/µl stock (ThermoFisher, 21580) in GlycoBuffer) twice for 15 min each at RT protected from light. The slides were washed once in 1X CODEX Buffer. A cotton-tipped applicator was used to scrape off the hydrophobic barrier on the slides and the slides were stored in 1X CODEX Buffer. Details regarding the lectins, antibody clones, vendors, conjugated channels, titers, exposure times, and assigned channels throughout the study are in **Supplementary Table 2**.

#### Imaging

GlycoScope imaging was performed via the automated PhenoCycler Fusion platform (Akoya Biosciences). Each tissue slide was mounted with a flow cell (Akoya Biosciences, 240205) and was incubated in 1X CODEX Buffer for 10 min at RT. A reporter plate was prepared for each tissue slide with each well corresponding to each imaging cycle. Briefly, each well of a 96-well black reporter plate (BRAND Tech, 781607) was added with Plate Buffer (500 µg/ml sheared salmon sperm DNA in 1x CODEX Buffer) supplemented with 54.11 mM Hoechst 33342 (ThermoFisher, H3570) and 100 nM each of ATTO550 or AlexaFluor647-conjugated lectin/antibody corresponding complementary reporter oligos (GenScript, HPLC purified). The wells were then sealed using aluminum plate seal (ThermoFisher, AB0626) and stored at 4°C until use. Low DMSO (80% 1× CODEX Buffer, 20% DMSO) and High DMSO (10% 1× CODEX Buffer, 90% DMSO) buffers were prepared fresh by mixing 1× CODEX Buffer with DMSO (Sigma, 472301-4L). After imaging, the slides were stored in 1× PBS.

### Hematoxylin & Eosin staining and imaging

All slides were stored in 1x CODEX buffer at 4°C after GlycoScope capture for H&E staining to visualize and confirm tissue morphology immediately after completing quality control evaluation of the captured images. Slides were first equilibrated in UltraPure water (Invitrogen 10977-023) at RT prior to staining with Modified Mayer’s Haematoxylin (StatLab, HXMMHPT) for 5 min at RT, followed by rinsing 3 times with UltraPure water (Invitrogen 10977-023). Slides were then treated with Bluing Solution (StatLab, HXB00588E) for 1 min at RT, and subsequently rinsed 3 times with UltraPure water (Invitrogen 10977-023). The slides were then equilibrated in 95% ethanol for 1 min prior to staining with a solution of Eosin Y and Phloxine B (StatLab, STE0243) for 1 min, followed by rinsing by dipping 12 times each in three changes of fresh 95% ethanol. Finally, the slides underwent graded dehydration by dipping 5 times in 100% ethanol, and 5 times in two changes of xylenes. Excess xylenes was gently dabbed off and glass coverslips (Creative Waste Solutions, CSM-2450) were mounted with xy-lene-based mounting medium (OptiClear Xylene, SSN Solutions, CSM1112). The slides were left to dry overnight at RT, after which they were scanned using the Grundium Ocus40 slidescanner (Grundium, MGU-00003). The H&E stains were verified by board certified pathologist W.R.B for tissue quality and morphological consistency with the multiplexed spatial proteomics images.

### Data analysis: GlycoScope

#### Image processing

For all markers in the tonsil dataset, images were derived from the 8-bit final QPTIFF files processed by the Pheno-cycler Fusion software with default blank cycles subtraction. These marker images were then extracted and subsequently subjected to cell segmentation and single-cell feature extraction, as described in the following sections.

For all antibody markers in the FL/FH dataset, the 16-bit intermediate QPTIFF files generated by the Phenocycler Fusion were used to ensure optimal dynamic range of data. The QPTIFFs were first processed by subtracting the last blank cycle scaled by the ratio between current channel cycle and total cycle number, i.e.,

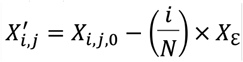

where X’_i,j_ is the blank-subtracted image of marker *j* in cycle *i*; X_i,j,0_ is the 16-bit intermediate image of marker *j* in cycle *i*; and Xε is the last blank cycle. These images were then processed in imageJ using the “Math” and “Subtract Background” functions under “Process” to 1) subtract the mean pixel value of the image to remove most of the “salt and pepper” noise, and 2) subtract the background generated by the sliding paraboloid algorithm with a 5 pixel radius. Finally, binary masks were generated for these images using Otsu binarization. Then the final images were generated by taking the Hadamard product of the binary masks and the background-subtracted images.

For all glycan markers (DC-SIGNL, MBL-L, Siglec-7L, Siglec-9L, Siglec-15L, Galectin-8L, Galectin-9L) in the FL/FH dataset, the 16-bit intermediate QPTIFF files generated by the Phenocycler Fusion were used and underwent the aforementioned image processing pipeline. In addition, these images were processed by nuclear masking. Briefly, respective nuclear DAPI images of the maker cycles were used to generate binary masks. These masks were then applied to the marker images to remove any non-specific nuclear signals.

#### Cell segmentation and single cell feature extraction

For both the tonsil and FL/FH GlycoScope datasets, cell segmentation was performed using the MESMER model of DeepCell version 0.12.2^70,71^, with maxima_threshold set to 0.075 and interior_threshold set to 0.05. The nuclear channel input of MESMER was DAPI for both datasets. The membrane channel input of MESMER for the tonsil dataset was a summation of HLA-1 and Galectin-8L, while that for the FL/FH datasets was a summation of HLA-1, CD31, CD3, CD68 and podoplanin.

For each marker, the pixel value within the area of each cell (determined by the segmentation mask) was summed and then divided by the area of each cell, and the resulting cell-size scaled sum was set as the expression value for a given marker. Each cell from each tonsil or core was identified by unique cell label and tissue ID.

#### Data processing

Cell sizes that were below 0.001 (i.e., non-specific dots or tissue debris) and over 0.995 percentile (i.e., due to flipping of tissue during imaging) were first removed. All the markers of the FL/FH dataset were then scaled by their respective median nuclear DAPI signal of each tissue sample to adjust for different binding efficiency of markers. Lastly, a local min-max scaling within each tissue was applied to the phenotyping marker (Podoplanin, Pax5, HLA-1, FoxP3, CD8, CD68, CD4, CD31, CD3, CD23, CD21, CD206, CD20, CD163, CD11c, CD11b, BCL6, BCL2, CD56); while a global min-max scaling was applied to the functional and glycan markers (Functional: Tox1/2, TIM3, TCF1, PD1, LAG3, Ki67, IRF4, HLA-DR, CD79A, CD45RO, CD45RA, CD40L, CD30, CD138, CD123; Glycans: DC-SIGNL, MBL-L, Siglec-7L, Siglec-9L, Siglec-15L, Galectin-8L, Galectin-9L), to scale the marker expression levels to be within [0,1].

#### Cell phenotyping

Cell phenotyping was performed on the processed FL/FH dataset through iterative rounds of clustering and annotating processes with PhenoGraph based on the phenotyping marker enrichment profiles (26095251). Clustering results were plotted as heatmaps showing the Z-score of each marker within each cluster and were annotated based on the combination of markers that had the highest Z-score in that cluster. Pax5, CD3, CD68, CD31, Podoplanin and CD21 were used to annotate the general cell types of B/tumor, T, myeloid, endothelial, lymphatic and FDC-enriched cells, respectively. Cells that showed unclear marker enrichment patterns were annotated as “Other” cells. B/tumor cells were further annotated into Gal8L^Low^ B, Gal8L^High^ B, Gal8L^High-^ BCL2+ B, Gal8L^High^BCL6+ B and Gal8^High^BCL2+BCL6+ B with the addition of BCL2, BCL6 and Galectin-8L markers. T cells were further annotated into CD4+ T, CD8+ T and Tregs with the addition of CD4, CD8 and FoxP3 markers. Myeloid cells were further annotated into M1-like (CD68+CD163-/CD206-) macrophages and M2-like (CD68+CD163+/CD206+) macrophages with the addition of CD163 and CD206 markers. After an initial round of clustering with PhenoGraph was performed, clusters with clear enrichment patterns were annotated, while clusters with mixed patterns underwent additional rounds of iterative clustering and annotation until all identifiable cells were annotated. To visualize and confirm the assigned annotations, Mantis Viewer^72^ was used to overlay the annotations onto the segmentation mask and the marker image, for visual inspection. The final annotations were verified by a board-certified pathologist W.R.B..

#### Coefficient of variation (CV) score

The coefficient of variation (CV) was used to assess the relative variability of each marker across the three experimental conditions: Antibody-only, Lectin-only, and GlycoScope. The CV is defined as:

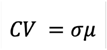

where σ is the standard deviation and µ is the mean expression of a given marker within each condition. CV scores were standardized with Z-score transformation and plotted as heatmap to facilitate comparison across markers and conditions. Heatmaps were plotted with R package ggplot2.

#### Marker enrichment heatmap

The marker enrichment heatmap showed the Z-score of a given (marker, cell type, disease state) tuple. In other words, it showed how many standard deviations away is the mean of marker A expression of cell type B given a disease state from the population mean of marker A expression:

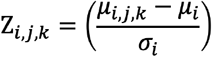

where Z_*i,j,k*_ stands for the Z-score for marker *i*, cell type *j* and disease status *k*; µ_i,j,k_ stands for the mean expression for marker *i*, cell type *j* and disease status *k*; µ_i_ stands for the population mean of marker *i*, and σ_i_ stands for the population standard deviation of marker *i*. Heatmaps were plotted with R package ggplot2.

#### Relative proportion of cell type

Relative proportion of cell type was presented as the ratio between the total number of cells within a given cell type with the total number of cells within the tissue sample of a disease state (or step):

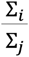

where ∑_*i*_ stands for the total number of cells in cell type *i*, and ∑_*j*_ stands for the total number of cells within the tissue sample (or step) *j*. Statistical analyses were performed with Kruskal-Wallis rank sum test followed by pairwise Wilcoxon test with Benjamini-Hochberg adjustments. Violin plots were plotted with R package ggplot2.

#### Mean expression of markers comparison

Mean expression of glycans or proteins across disease states and cell type were presented as the mean pixel intensity of each of the glycans or proteins in a given disease state and cell type at single cell level, respectively:

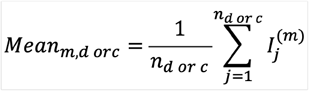

where *Mean*_*m,d or c*_ stands for mean expression of glycan or protein *m* in disease state *d* or cell type *c, I* ^(m)^_*j*_ stands for the mean of glycan or protein in cell *j*, and *n*_*d or c*_ stands for the number of cells in disease state *d* or cell type *c*. Statistical analyses were performed with Kruskal-Wallis rank sum test followed by pairwise Wilcoxon test with Benjamini-Hochberg adjustments. Violin plots were plotted with R package ggplot2.

#### Follicular dendritic cell (FDC)-anchored spatial gradient

For each tissue core, Euclidean distances from the centroids of follicular B cells, including cells from the mantle zone and follicular center, to the nearest FDC were computed using k-nearest neighbor search. Analyses were performed independently within each core to account for core-level variability. To visualize distance-dependent patterns, DC-SIGNL expression in follicular B cells was summarized at the core level across predefined distance bins from the nearest FDC (0-10, 10-20, 20-40, 40-60, 60-100, and 100-150 µm), and disease-level means with 95% confidence intervals were calculated from these core-level summaries. Line graphs were plotted with R package ggplot2.

#### Spatial analysis anchoring from mantle zone rim

Binary masks delineating the Galectin-8L–expressing B cells at the mantle zone rim were initially generated using ImageJ. These masks were then used to create concentric expansion zones both inwards and outwards at 10 µm intervals, which were overlaid onto the cell segmentation mask. Each cell falling within a given zone was assigned a corresponding step number, enabling analysis of relative cell type proportions and mean marker expression as described above. Bar plots and line graphs were plotted with R package ggplot2.

#### Correlation analysis

Spearman’s rank correlation coefficients between the proportion of markers expressing T cells (i.e., PD1, Tox1/2, TIM3, LAG3, CD3) and Galectin-8L (or BCL2) mean expression in B cells at each defined step within the follicle were calculated. All p-values were adjusted for multiple comparisons using the Benjamini-Hochberg method. Correlation scatter plots were plotted with R package ggplot2.

---

**Fig S1.**
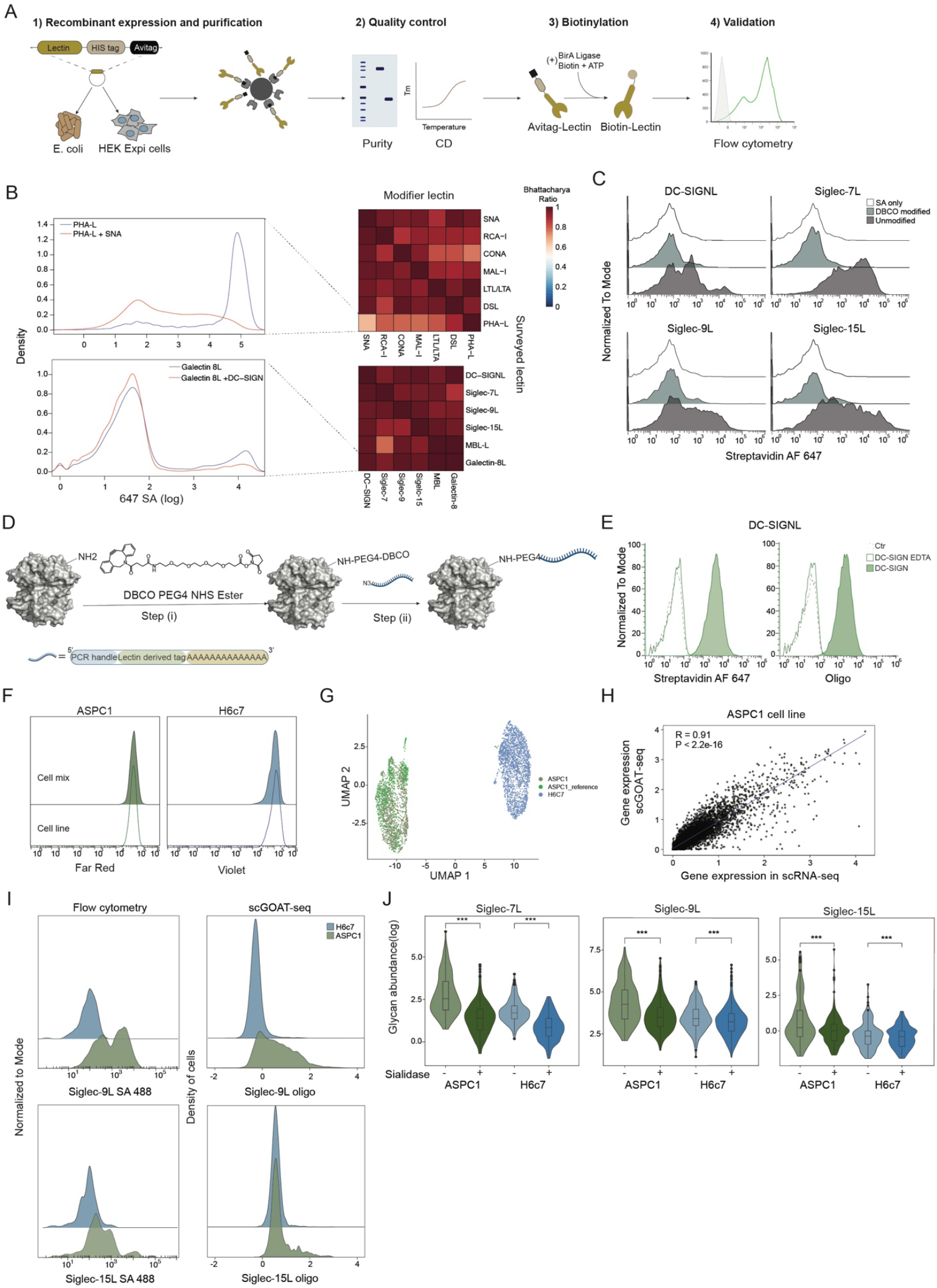
Validation and benchmarking of human recombinant lectins and scGOAT-seq assay. **(A)** Overview of human recombinant lectin production and validation. (**B**) PBMCs were stained with either single recombinant human lectins or equimolar lectin pairs, and binding distributions were compared using the Bhattacharyya overlap coefficient. Left, representative smoothed density histograms show kernel-density estimates of fluorescence intensities, illustrating the underlying distribution of lectin-binding. Right, heatmap shows averaged overlap values (n = 3; 1 = identical distributions). **(C)** Representative flow cytometry histograms showing PBMC staining with each lectin before and after NHS-DBCO modification. **(D)** Schematic representation of streptavidin single-strand oligonucleotide labeling, incorporating an oligo-dT primer, a lectin-derived tag, and a PCR amplification handle. **(E)** Representative histograms of flow cytometry analysis comparing oligoor fluorescently labeled DC-SIGN binding, with EDTA as a negative control. **(F)** Representative histograms of flow cytometry analysis of the artificial pancreatic tumor model and individual pancreatic cell lines (ASPC1 and H6c7) after labeling with cell tracer dye. **(G)** UMAP visualization of cells, colored by cell line and reference map. **(H)** Genegene correlation analysis comparing scRNA-seq and scGOAT-seq for ASPC1 cells. **(I)** Representative histograms showing Siglec-9L and Siglec-15L abundance per cell line in the cell mix model analyzed by both, flow cytometry and scGOAT-seq. (**J**) Violin plot showing Siglec-7L, Siglec-9L and Siglec-15L abundance across cell types in the cell mix model under basal conditions or after sialidase treatment (Wilcoxon rank-sum test). ***P<0.001.

**Fig S2.**
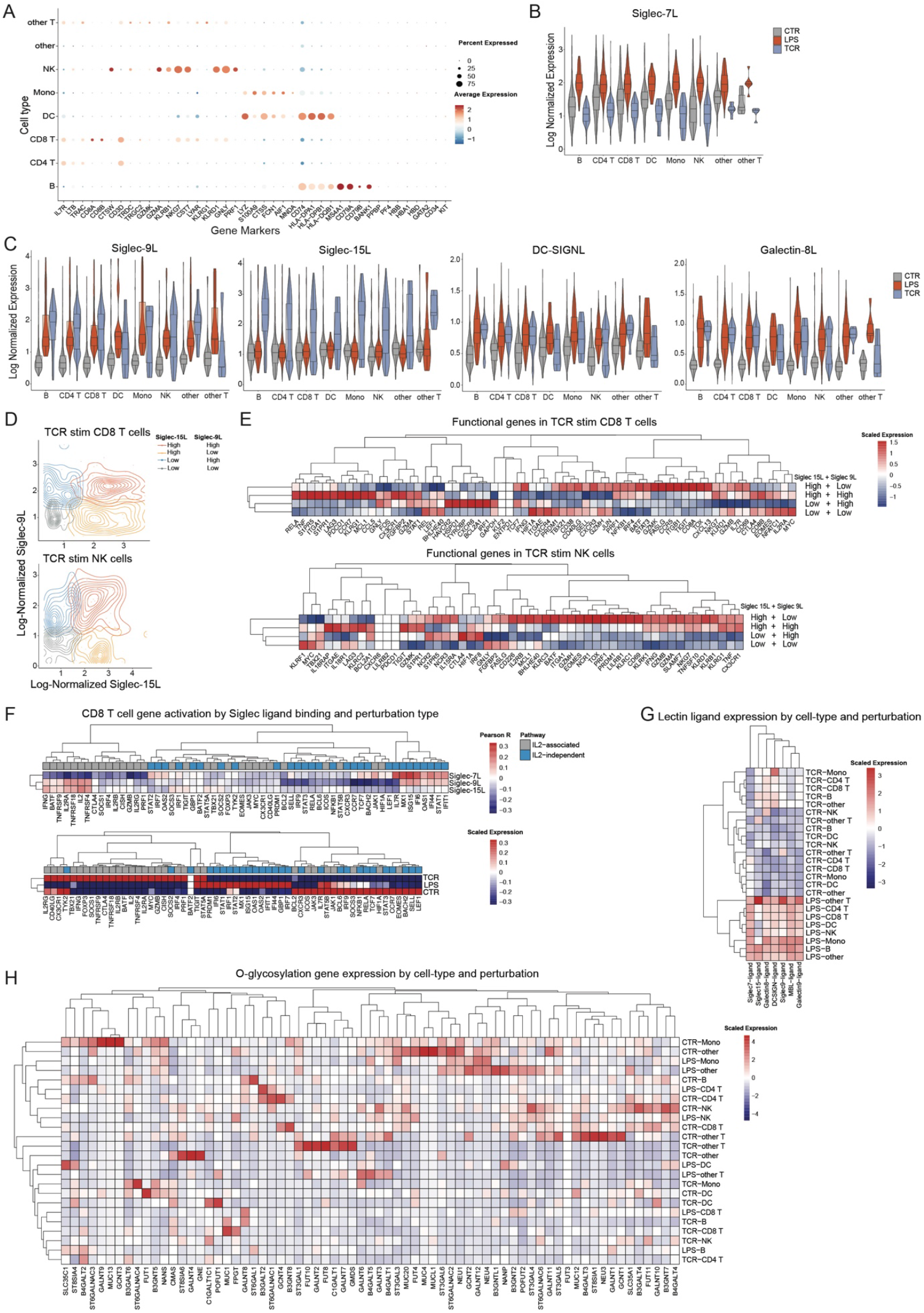
scGOAT-seq identifies novel T cell glyco-states associated with immune perturbations. **(A)** Dot plot showing the expression of different marker genes used for cell type annotation across different cell types. **(B-C)** Violin plots depicting log-normalized expression of lectin-binding ligands for Siglec-7L, Siglec-9L, Siglec-15L, DC-SIGNL, and Galectin-8L across cell types and perturbations. **(D)** Scatter density plot showing different populations of TCR stimulated CD8+ T cells and NK cells binned by expression levels of Siglec-9L and Siglec-15L. **(E)** Heatmap of key functional CD8+ T cell and NK cell gene expression across TCR stimulated CD8+ T cells and NK cells grouped by expression levels of Siglec-9L and Siglec-15L. **(F)** Heatmap showing Pearson correlation of functional CD8+ T cell activation genes with Siglec-7L, Siglec9L and Siglec-15L (top) and their expression across CD8+ T cells in different perturbations (bottom). **(G)** Expression and clustering of different lectin-ligands across different cell types and conditions. **(H)** Expression and clustering of O-glycosylation related genes across different cell types and conditions.

**Fig S3.**
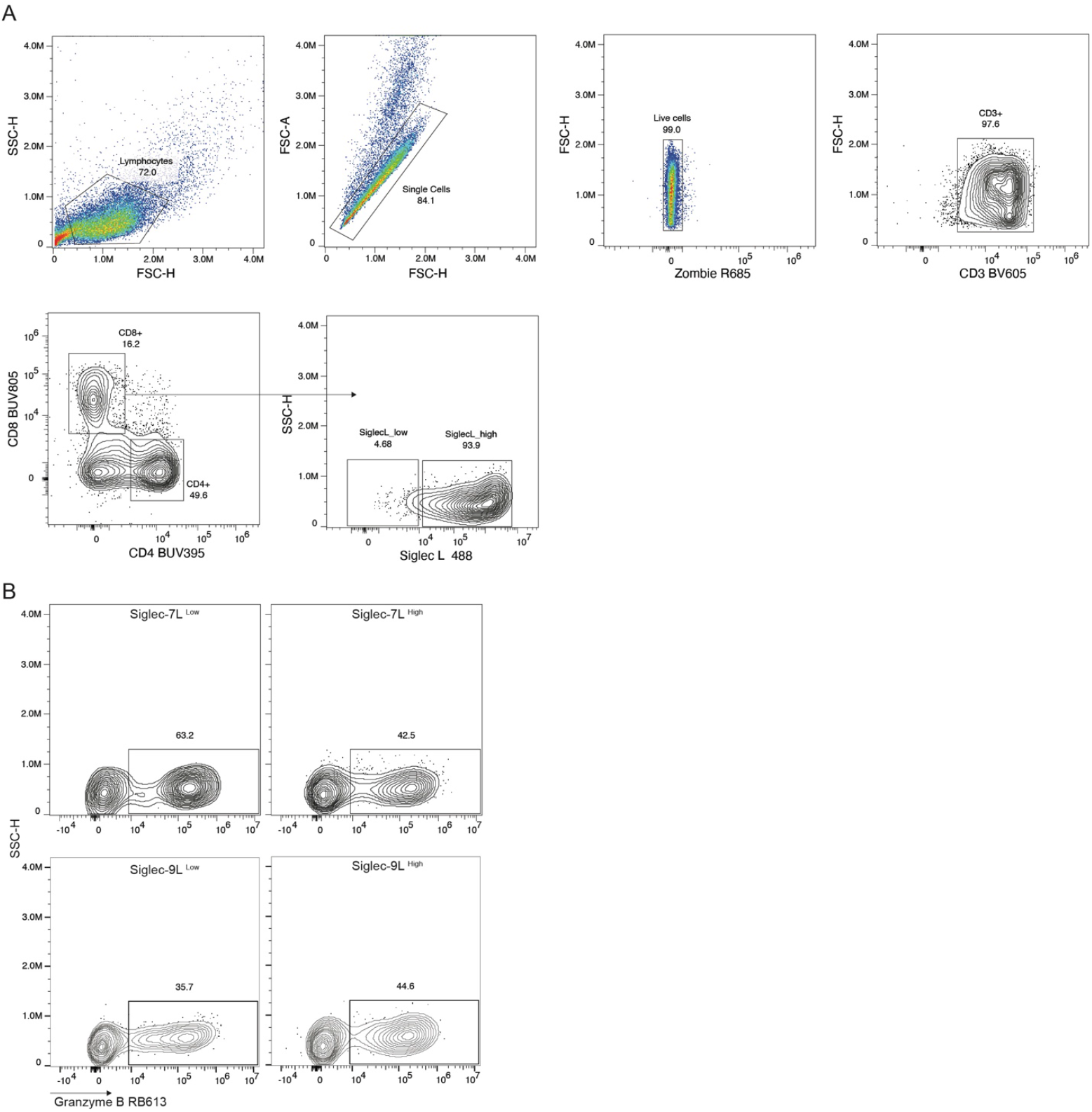
Gating strategy and Siglec-ligand–based stratification of perforin and granzyme B expression in TCR-activated CD8^+^ T cells. **(A)** Representative spectral flow cytometry gating strategy. Events were first gated by FSC-H and SSC-H to exclude debris, followed by singlet discrimination (FSC-A vs SSC-H). Live cells were identified using Zombie viability dye, and CD3^+^ T cells were subsequently gated and subdivided into CD8^+^ and CD4^+^ populations. Within the CD8^+^ compartment, cells were stratified into Siglec-L^High^ and Siglec-L^Low^ populations based on probe binding intensity. **(B)** Spectral flow cytometry gating of CD8+ T cells after TCR activation, with independent stratification by Siglec-7 or Siglec-9 ligand binding and assessment of granzyme B expression within each Siglec-defined population.

**Fig S4.**
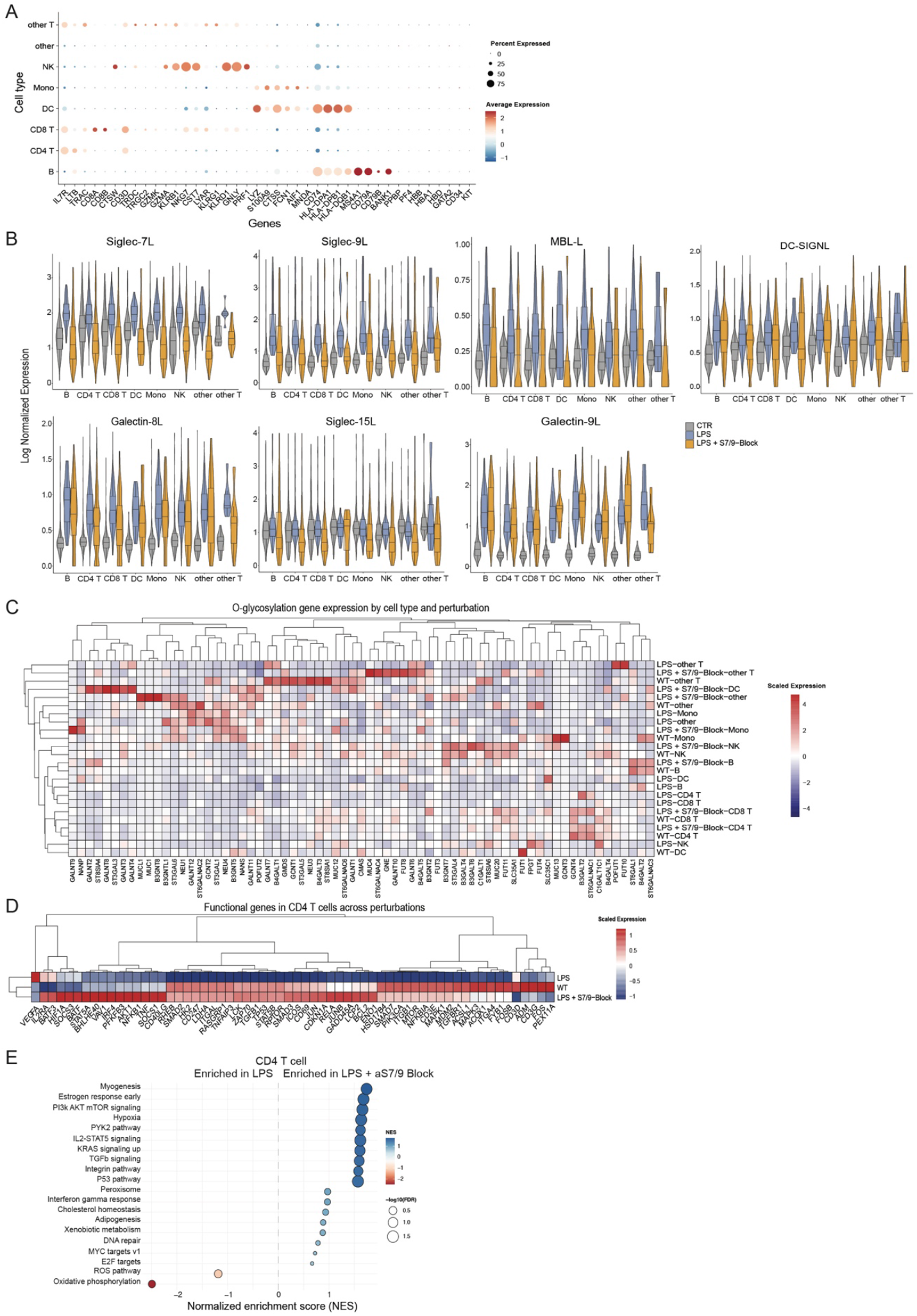
scGOAT-seq identifies immune activation changes associated with blocking Siglec-mediated glycan recognition. **(A)** Dot plot showing the expression of different marker genes used for cell type annotation across different cell types. **(B)** Violin plots depicting log-normalized expression of lectin-binding ligands for DC-SIGNL, Galectin-8L, MBL-L and Siglec-15L across cell types and perturbations. **(C)** Expression and clustering of O-glycosylation related genes across different cell types and conditions. **(D)** Heatmap of key functional CD4+ T gene expression across perturbations. **(E)** GSEA analyses across CD4+ T between LPS stimulation with anti-Siglec7 and anti-Siglec9 blocking antibodies compared to LPS treatment alone.

**Fig S5.**
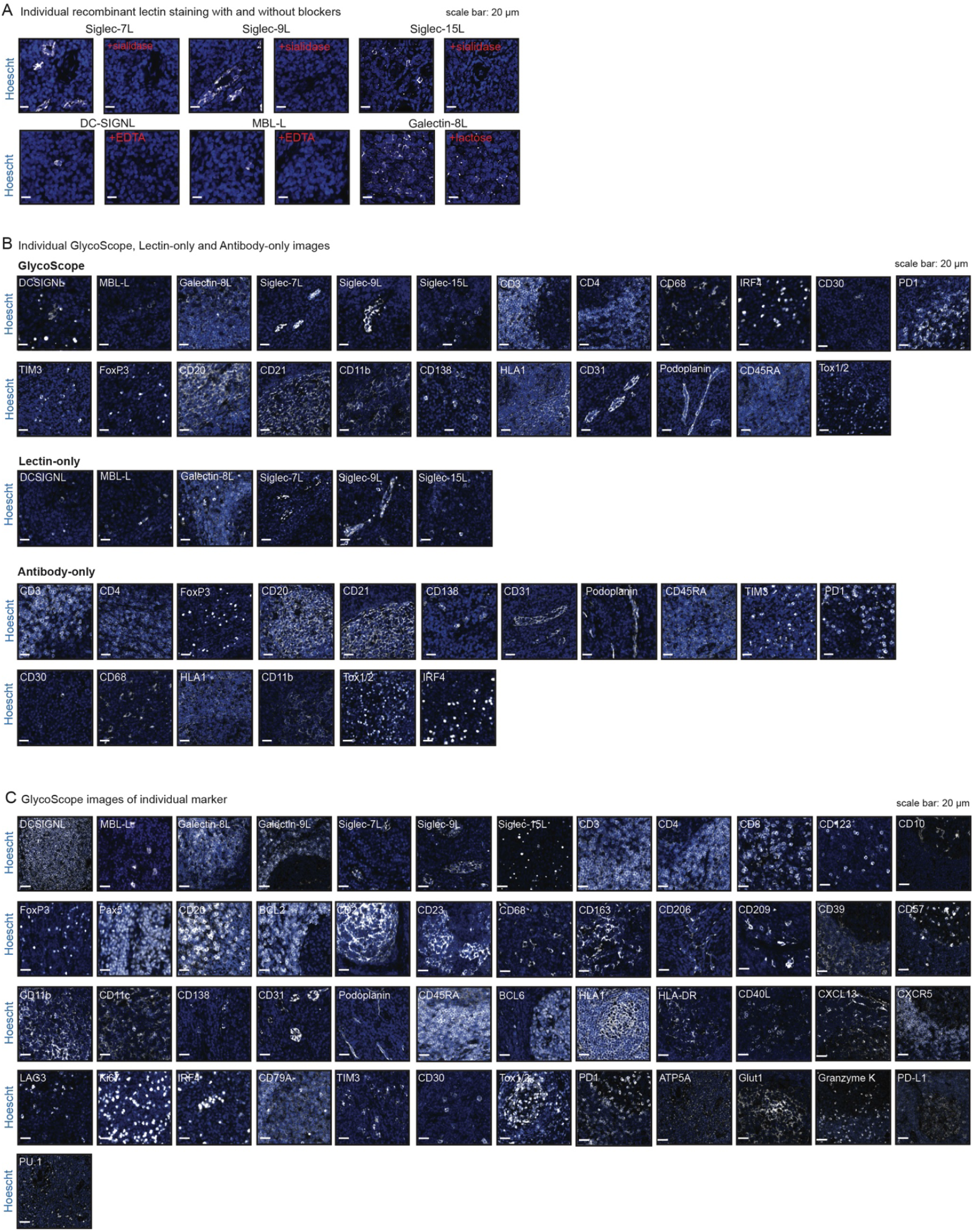
Validation and benchmarking of GlycoScope. **(A)** Representative multiplexed images showing nuclear staining (DAPI, blue) and individual glycans (white) with and without their corresponding blockers (scale bar: 10 µm). **(B)** Representative individual images of 24-plex lectin-antibody panel from benchmarking study across GlycoScope, Lectin-only and Antibody-only conditions (scale bar: 20µm) shown for subcellular staining specificity purposes. **(C)** Representative individual images of 32-plex lectin-antibody panel from the FL study acquired from GlycoScope (scale bar: 20µm) shown for subcellular staining specificity purposes.

**Fig S6.**
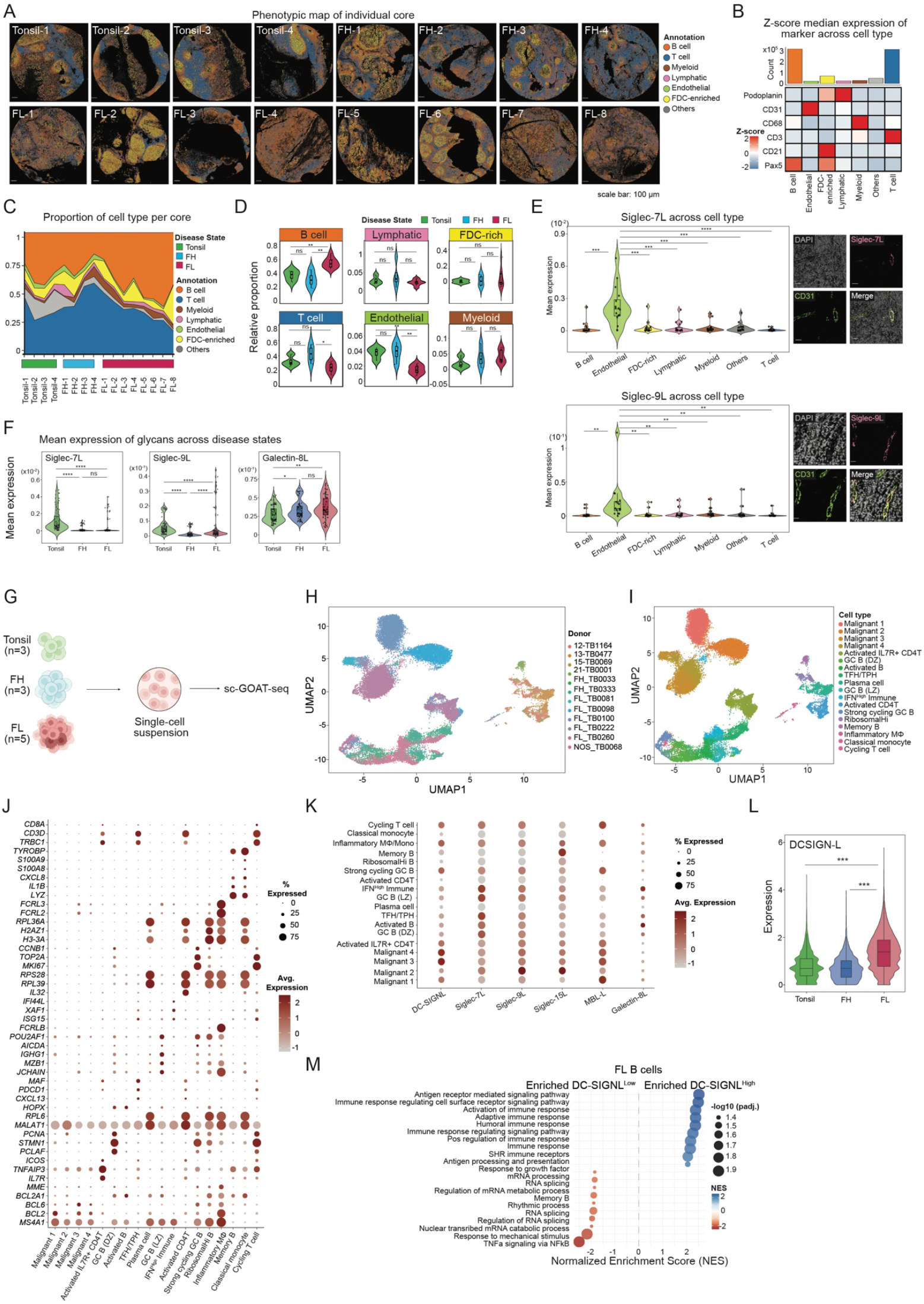
Integrated spatial and single-cell glycomics recapitulate known mannosylated-BCR biology in follicular lymphoma. **(A)** Phenotype maps of tonsil, FH and FL tissue sample cores (scale bar: 100µm). **(B)** Heatmap showing median expression of protein markers across cell type: B (Pax5+), endothelial (CD31+), FDC-enriched (Pax5+ CD21+ Podoplanin+), lymphatic (Podoplanin+) and T cells (CD3+). **(C)** Proportion of cell types across all cores. **(D)** Relative proportion of cell types across FL, FH and tonsil tissues (Kruskal-Wallis rank sum test followed by pairwise Wilcoxon test with Benjamini-Hochberg adjustments). *P<0.05; **P<0.01; ****P<0.0001; ns P>0.05. **(E)** Violin plot showing the mean expression of Siglec-7L (left) and Siglec-9L (right) across cell types (Kruskal-Wallis rank sum test followed by pairwise Wilcoxon test with Benjamini-Hochberg adjustments). **(F)** Mean expression of glycans across tonsil, FH and FL tissues. (Kruskal-Wallis rank sum test followed by pairwise Wilcoxon test with Benjamini-Hochberg adjustments). **(H)** UMAP representation of donors from FH, FL and tonsil samples. **(I)** UMAP representation of cell types from follicular lymphoma samples. **(J)** Average expression of cell type-specific marker genes across each cell population identified by scGOAT-seq. **(K)** Average expression of glycans in each individual cell type identified from scGOAT-seq. **(L)** Violin plots showing the abundance of DC-SIGNL in tonsil, FH and FL cells acquired from scGOAT-seq (Pairwise Wilcoxon test). **(M)** Gene set enrichment analysis comparing DC-SIGNL^Low^ and DC-SIGNL^High^ FL B cells. NES: Normalized Enrichment Score. ***P<0.001; ****P<0.0001.

**Fig S7.**
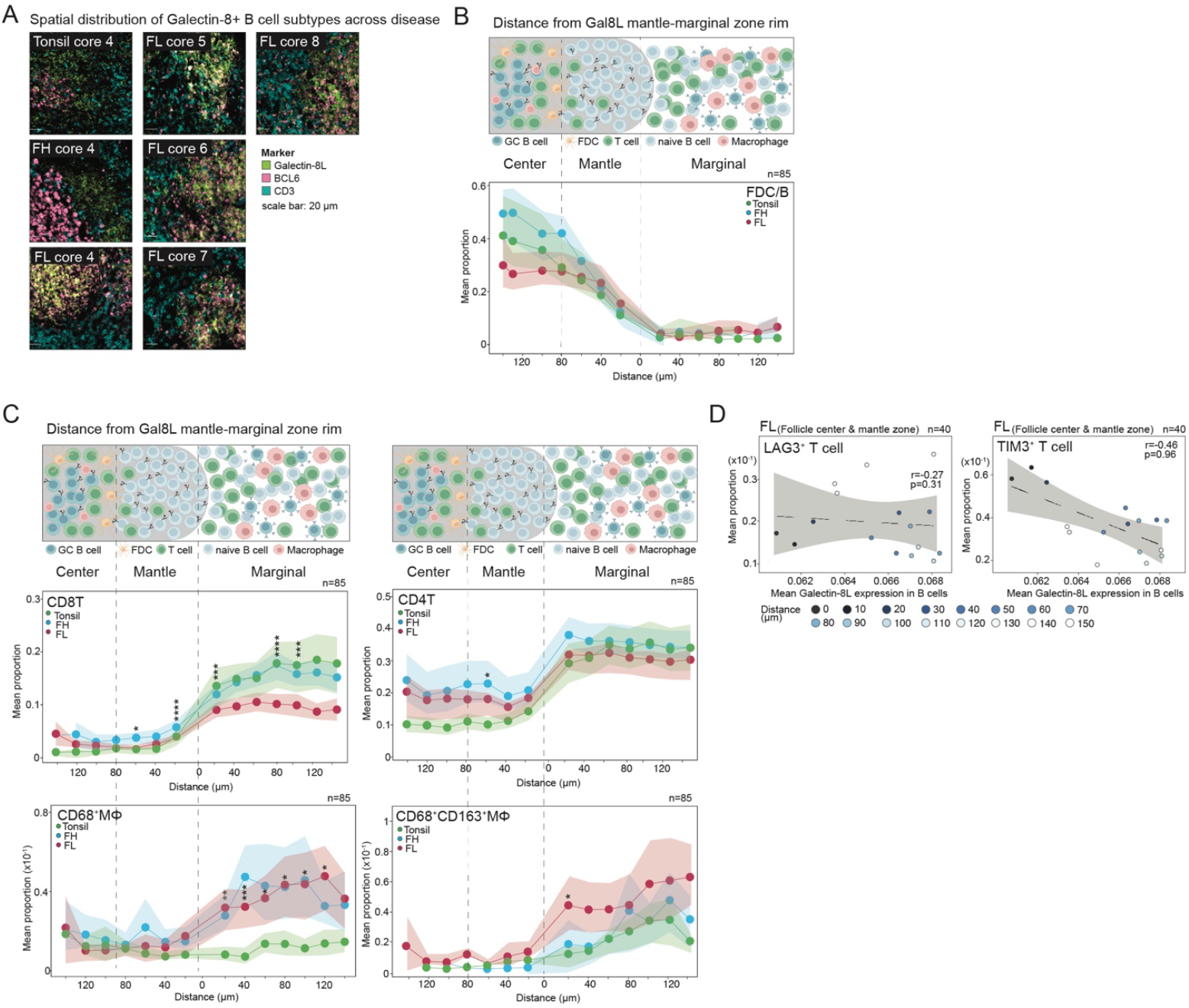
GlycoScope maps a distinct Galectin-8L spatial pattern in FL neoplastic follicles that associate with lectin proximity and distinct intrafollicular immune niches. **(A)** Additional multiplexed images of tonsil, FH and FL cores showing Galectin-8L (green), BCL6 (pink) and CD3 (cyan) markers (scale bar: 20µm). **(B)** Mean proportions of FDC-enriched B cells between 0-140µm towards the follicle center and marginal zone from the anchor rim are shown (Kruskal-Wallis rank sum test followed by pairwise Wilcoxon test with Benjamini-Hochberg adjustments). **(C)** Mean proportions of CD8+ T (top left), CD4+ T (top right), CD68+ Macrophage (bottom, left) and CD68+ CD163+Macrophage (bottom right), between 0-140µm towards the follicle center and marginal zone from the anchor rim are shown (Kruskal-Wallis rank sum test followed by pairwise Wilcoxon test with Benjamini-Hochberg adjustments). **(D)** Correlation scatter plot of the proportion of LAG3+ (left) and TIM3+ (right) T cells with Galectin-8L expression in B cells at each step within the follicle (Spearman’s correlation with Benjamini-Hochberg adjustments). *P<0.05; **P<0.01; ****P<0.0001; ns P>0.05.

## Notes

### Competing Interest Statement

The authors have declared no competing interest.

